# FGF diffusion is required for directed migration of postembryonic muscle progenitors in *C. elegans*

**DOI:** 10.1101/2025.03.19.643693

**Authors:** Theresa V Gibney, Ariel M Pani

**Affiliations:** Department of Biology, University of Virginia, Charlottesville, VA USA; Department of Cell Biology, University of Virginia School of Medicine, Charlottesville, VA USA

**Keywords:** Fibroblast Growth Factor, FGF, diffusion, gradient, cell migration

## Abstract

Extracellular signaling molecules mediate crucial aspects of cell-cell communication and play essential roles in development and homeostasis. Fibroblast Growth Factors (FGFs) are a conserved family of secreted signaling proteins that can disperse long distances between cells and are often thought to form concentration gradients that encode spatial information. However, we know relatively little about the spatial distribution of FGFs *in vivo*, and endogenously tagged FGFs use different mechanisms to move between cells and form gradients in zebrafish and flies. We used FGF-dependent migration of *C. elegans* muscle progenitors called sex myoblasts (SMs) as a tractable system to elucidate FGF dispersal mechanisms and dissect how FGF guides migrating cells. Live imaging of cell dynamics and endogenously tagged FGF combined with membrane tethering and extracellular trapping approaches revealed that endogenous FGF is diffusible *in vivo* and extracellular dispersal is required for SM migration. Misexpression demonstrated FGF is a bona fide chemoattractant that orients SMs during a critical window, while an unidentified, short-range signal acts in concert to precisely position SMs. Our finding that an invertebrate FGF is endogenously diffusible suggests that this may be the ancestral mode for FGF dispersal.

**Summary Statement:** Fibroblast Growth Factors (FGFs) are signaling proteins with crucial developmental roles. We demonstrated that a *C. elegans* FGF homolog is diffusible, and diffusion is required *in vivo* for cell migration guidance.

## Introduction

Cell-cell signaling and dynamic cellular interactions orchestrate animal development, homeostasis, and regeneration. During development, cells use secreted signaling proteins to communicate at a range of distances to regulate patterning, growth, differentiation, and cell behaviors such as migration and morphogenesis (Wolpert, 2016, Muller and Schier, 2011). How secreted signaling proteins move between cells to reach their destinations *in vivo* remains largely unknown. Fibroblast Growth Factors (FGFs) are a family of secreted signaling proteins with essential roles in animal development (Dorey and Amaya, 2010, Thisse and Thisse, 2005), and abnormal FGF signaling is implicated in disease states (Turner and Grose, 2010, Babina and Turner, 2017). FGF proteins are often thought to form concentration gradients that encode spatial information and act as morphogens that regulate patterning and cellular behaviors over long distances (Dorey and Amaya, 2010, Muller et al., 2013, Thisse and Thisse, 2005, Balasubramanian and Zhang, 2016). Yet, endogenous FGF protein gradients have only been observed in a handful of cases (Du et al., 2018, Harish et al., 2023, Toyoda et al., 2010, Dubrulle and Pourquie, 2004, Chen et al., 2009), and how endogenous FGF proteins move between cells *in vivo* has been debated based on data from a limited number of models. In vertebrates, endogenously tagged FGF8a forms long-range gradients by diffusion in developing zebrafish (Harish et al., 2023), and *fgf8* mRNA decay coupled with growth contribute to an anteroposterior protein gradient in mouse and chick embryos (Dubrulle and Pourquie, 2004). However, two *Drosophila* FGFs are not freely diffusible and instead signal primarily at cell contacts (Stepanik et al., 2020, Du et al., 2022, Du et al., 2018). In the absence of extracellular diffusion, FGFs can move long distances between cells using dynamic cell membrane extensions known as cytonemes that physically link signaling and responding cells (Du et al., 2022, Du et al., 2018).

Major challenges for investigating how FGFs disperse in living animals include difficulties with visualizing endogenous signaling proteins and their hypothesized gradients *in vivo* along with challenges observing cytonemes, which are not preserved by standard fixation methods or visible using cytosolic fluorescent proteins (Sanders et al., 2013, Kornberg, 2017). Studies of FGF diffusion and gradient formation have often relied on exogenous fusion proteins, with the caveats that transgenic approaches often do not recapitulate native expression patterns, and overexpression may alter extracellular dispersal dynamics and/or cell behaviors. Of the endogenously tagged FGF proteins examined so far, zebrafish FGF8a and *Drosophila* Branchless have been found to disperse between cells using different mechanisms (Du et al., 2022, Harish et al., 2023, Du et al., 2018), and data from additional models are needed to discern underlying principles.

An ideal system to investigate FGF dispersal would allow for live imaging of FGF-secreting and FGF-responding cells, visualization of endogenous FGF protein *in vivo,* and the ability to manipulate endogenous FGF dispersal without disrupting protein function or cell physiology. The migration of two *C. elegans* muscle progenitors, known as sex myoblasts (SMs), provides an experimentally tractable, *in vivo* model to investigate FGF signaling mechanisms. All *C. elegans* postembryonic mesoderm cells, including SMs, are derived from a single M mesoblast cell, which is located near the tail in early larvae. The M lineage gives rise to body wall muscles, coelomocytes, and two SMs that are located on the left and right sides of the animal (Sulston and Horvitz, 1977) (see Fig. 1). During the L2 larval stage, SMs migrate anteriorly to a precise position at the center of the gonad under the control of the FGF8/17/18 family homolog EGL-17 (Burdine et al., 1998, Burdine et al., 1997, Stern and Horvitz, 1991) and the FGF Receptor (FGFR) EGL-15 (DeVore et al., 1995, Lo et al., 2008, Stern and Horvitz, 1991). The prevailing model for SM migration is that FGF/EGL-17 is produced by the P6.p 1° vulval precursor cell (VPC) and uterine cells and acts as a chemoattractant that guides myoblasts to their final positions (Branda and Stern, 2000, Burdine et al., 1998, Sherwood and Plastino, 2018).

**Figure 1.**
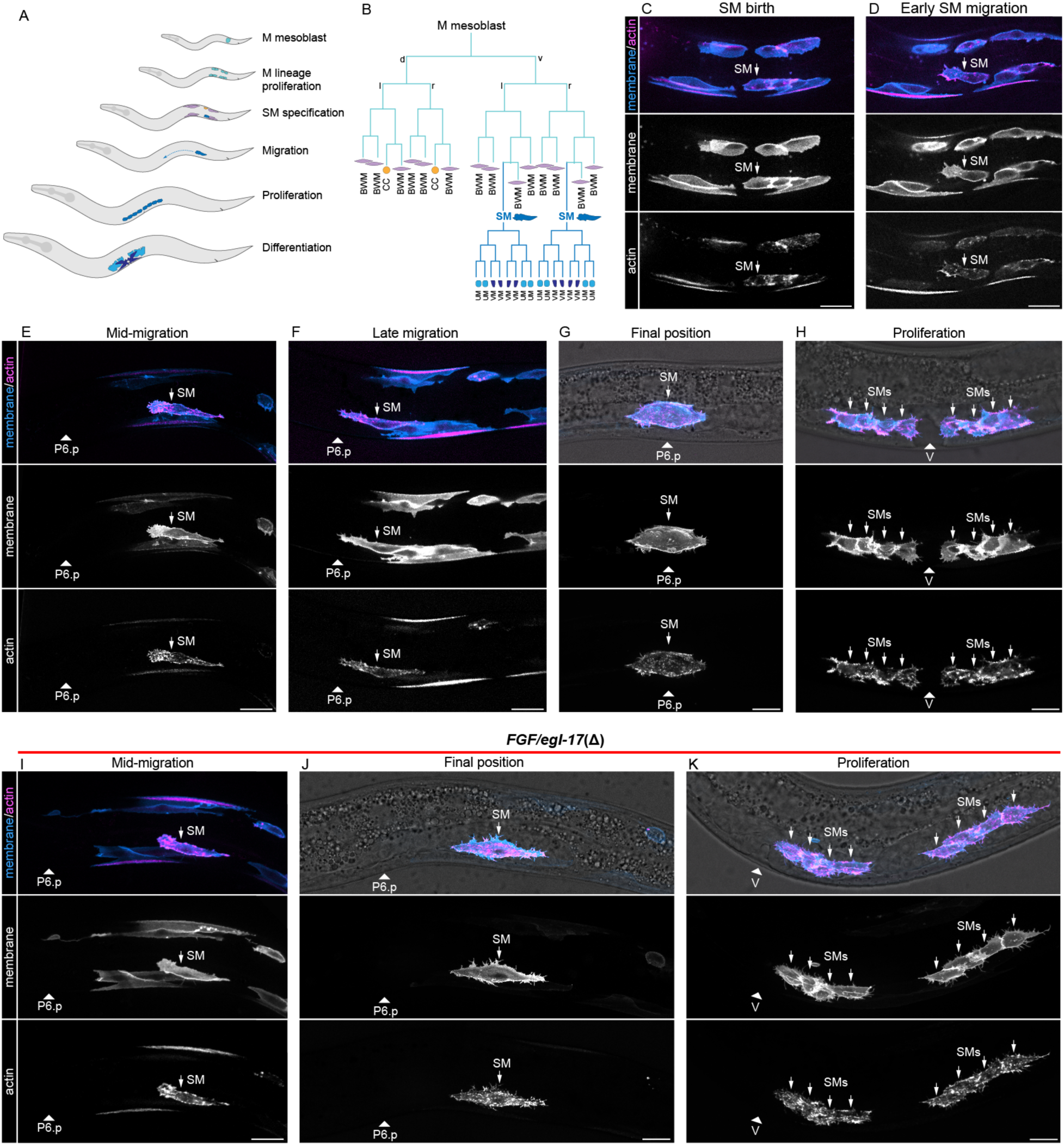
*In vivo* imaging of SM migration. (**A**) Schematic diagram of key developmental processes in the postembryonic mesoderm. (**B**) The M mesoblast gives rise to body wall muscles, coelomocytes, and sex myoblasts (SMs). Sex myoblasts migrate anteriorly to flank the center of the gonad, proliferate, and differentiate into vulval and uterine muscles. (**C**-**H**) *in vivo* imaging of M lineage and SM cell dynamics using a single copy *Phlh-8>2x mKate2::moesin ABD::F2A::2x mTurquiose2::PH* transgene to visualize plasma membranes (blue) and actin (magenta). (**D**) At the onset of SM migration, SMs delaminate from their sister body wall muscle cells and migrate anteriorly with broad, actin rich protrusions and short filopodia. (**E**, **F**) SMs exhibit similar protrusive behaviors as they crawl towards, and over the M-derived v17 and v18 body wall muscles towards the gonad center. (**G**, **H**) At the end of their migration, SMs center over the uterine cells, dorsal to the 1° VPC P6.p (**G**), before proliferating to form two clusters of four cells flanking the vulva (**H**). (**I**-**K**) In *FGF/egl-17(Δ)* animals, SM migration (**I**) is initially indistinguishable from wild-type, but SMs do not maintain polarized protrusions and fail to migrate past v17/18 (**J**). (**K**) SMs then proliferate in a posteriorly displaced location, leading to non-functional egg-laying muscles. All animals oriented with anterior to left and dorsal to top. Arrowheads denote the position of P6.p or the vulva depending on developmental stage. Abbreviations: BWM, body wall muscle; CC, coelomocyte; SM, sex myoblast; UM, uterine muscle; V, vulva; VM, vulval muscle. Autofluorescent gut granules were removed in E-K by subtracting a 445ex/642-80em background channel (see Fig. S3). Scale bars = 10 µm.

Here, we used live imaging and genome engineering to characterize interactions between SMs and *FGF/egl-17*-expressing cells, visualize endogenous FGF/EGL-17 distribution, and test roles for diffusion in SM migration. Live imaging of FGF-expressing cells and migrating SMs revealed SMs migrate in close proximity to previously unknown FGF source cells but are not linked through cytonemes. Intriguingly, endogenously tagged FGF did not form a visible protein gradient and localized primarily to FGF-expressing cells and SMs. Given the unexpected absence of a visible gradient, we used misexpression experiments to reassess the extent to which FGF acts as a permissive versus instructive signal in SM migration. Cell autonomous, spatially uniform, and posterior FGF misexpression phenotypes demonstrated that FGF orients migrating SMs towards an FGF source, but only during a critical time window. Anterior FGF misexpression also revealed somatic gonad cells can capture SMs migrating in close proximity, suggesting that an unidentified, short-range signal acts together with directional information encoded by FGF to mediate precise positioning of SMs. Finally, extracellular protein trapping approaches and an endogenously membrane-tethered FGF/EGL-17 demonstrated that endogenous FGF is diffusible, and its diffusion is required for SM migration.

## Results

### SMs migrate in close proximity to *FGF/egl-17*-expressing cells but do not interact through cytonemes

Distinguishing between potential mechanisms for secreted protein dispersal between cells requires knowledge of cellular shapes and behaviors to assess the possibility of direct contact between cells. To investigate FGF dispersal mechanisms during SM migration, we first characterized the architectures and behaviors of SMs throughout their migration. Although the *C. elegans* M lineage has been studied as a model for cell fate decisions and cell migration over several decades (Chen and Stern, 1998, Branda and Stern, 2000, Sherwood and Plastino, 2018, Liu and Murray, 2023), previous studies have focused largely on endpoint data, and high-resolution images of membrane and cytoskeletal architectures were not available. SMs are born in the L2 larval stage from an asymmetric division that gives rise to one SM and one body wall muscle (Sulston and Horvitz, 1977) (Fig. 1A, B). Using live imaging of plasma membrane and actin markers (Edwards et al., 1997), we observed that SMs delaminated from their sister body wall muscle cells and generated broad, actin-rich pseudopods polarized towards the anterior (Fig. 1C, D). SMs migrated anteriorly with sustained actin-rich protrusions at the leading edge of the cell, along with short filopodia. SMs migrated anteriorly for ∼20 um before contacting and then crawling over the dorsal surfaces of the M-lineage-derived ventral 17^th^ and 18^th^ body wall muscles (hereafter v17/18) (Fig 1E, F), which are visible with the same transgene used to visualize SMs. After migrating over the M-derived body wall muscle cells, SMs continued to produce broad protrusions at the leading edge as cells migrated over the surface of the somatic gonad, separated from underlying uterine cells by a basement membrane (Fig. S1). SMs then centered over the uterine progenitors (Fig. 1G), located dorsal to the primary vulval precursor cell (VPC) P6.p. After centering over the somatic gonad, each SM then underwent three rounds of division to generate two groups of four cells flanking the vulva (Fig. 1H). Throughout SM migration, we observed short filopodia, often near the leading edge of migrating cells, but did not observe cytoneme-like structures that could obviously link migrating SMs to the known FGF-expressing P6.p and uterine cells thought to produce the FGF/EGL-17 signal required for SM migration.

As a baseline to characterize phenotypes caused by manipulating FGF signaling, we deleted the entire *FGF/egl-17* coding region and characterized SM migration and actin dynamics. SM migration invariably failed in *FGF/egl-17(Δ)* animals, and the posteriorly displaced SMs in this strain resembled published data on SM migration in *FGF/egl-17* and *FGFR/egl-15(5a)* mutants. Live imaging in *FGF/egl-17(Δ)* animals showed that SM migration was unaffected until the point where SMs normally crawled over the v17/18 body wall muscles (Fig. 1I), which corresponds to the previously reported gonad- and FGF-independent early phase of SM migration (Branda and Stern, 2000). In *FGF/egl-17(Δ)* animals, after coming to rest atop the v17/18 body wall muscle cells, SMs failed to continue migrating (Fig. 1J) and no longer produced anteriorly polarized protrusions. SMs then proliferated in this posteriorly displaced location (Fig. 1K) leading to non-functional egg laying muscles. To evaluate the possibility that differential capacity for FGF-Ras-ERK signaling underlies the transition to FGF-dependent migration, we examined ERK-nKTR biosensor (de la Cova et al., 2017) activity in SMs during the FGF-independent and FGF-dependent stages of migration. We found that ERK-nKTR activity was not significantly different between FGF-independent and FGF-dependent time points (Fig. S2) indicating that changes in ERK activity are unlikely to underlie differential functions for FGF during early and late SM migration.

After failing to observe cytoneme-like structures in migrating SMs, we turned our attention to the identities and architectures of cells expressing *egl-17*, the FGF ligand required for SM migration. Transgenes driven by *egl-17* cis-regulatory sequences are expressed in the P6.p and dorsal uterine cells located at the endpoint of SM migration (Branda and Stern, 2000), which is key evidence supporting the model that FGF/EGL-17 acts as a chemoattractant to guide migrating SMs to the somatic gonad center. However, transgenes often do not recapitulate endogenous expression, and the native expression pattern and dynamics for *egl-17* was not known. To visualize *FGF/egl-17-*expressing cells *in vivo*, we used Cas9-triggered homologous recombination to engineer an endogenous bicistronic gene encoding *egl-17* along with a plasma membrane marker by inserting *SL2::mNeonGreen(mNG)::PH* at the 3’ end of the endogenous *egl-17* coding sequence after the stop codon (Fig. 2B). *egl-17::SL2::mNG::PH* homozygotes were phenotypically indistinguishable from wild-type and did not display SM migration defects. We used spinning disk confocal live imaging to characterize the *FGF/egl-17* expression pattern from the early L1 through L3 stages, focusing on the posterior half of the animal where the M lineage develops. To better visualize low levels of fluorescent protein expression in areas with autofluorescent gut granules, we removed broad-spectrum autofluorescence by subtracting a simultaneously acquired background channel (see Fig. S3). In young animals, prior to the first M cell division we observed *egl-17::SL2::mNG::PH* expression in Q neuroblasts along with weaker expression in the V5 seam cells and a rectal cell. We also observed variable and faint fluorescence in the M cell itself (Fig. 2C). *egl-17::SL2::mNG::PH* expression then shifted near the time of the first M cell division, unexpectedly becoming highly expressed in ventral midline blast cells (Fig. 1D) along with the M lineage cells themselves (Fig. 2D, E). As animals continued to develop, *egl-17::SL2::mNG::PH* expression diminished in M lineage cells but was maintained at high levels in a continuous row of cells along the ventral midline (Fig. 2F, G; Fig. S4). After the birth of the SM, we no longer observed *egl-17::SL2::mNG::PH* expression in the M lineage. During the early phase of SM migration, *egl-17::SL2::mNG::PH* remained highly expressed in the P4.p-P8.p blast cells, with lower expression in their neuronal progeny (Fig. 2H). We also observed low, but consistent expression in V5-derived neuroblasts located on the dorsal side of the animal, slightly posterior to the concurrent position of migrating SMs (Fig. 2I). The close association between *FGF/egl-17*-expressing cells and migrating SMs at this time was reminiscent of the association between *FGF/branchless-*expressing cells and growing tracheal branches in *Drosophila* embryos (Du et al., 2017). As SMs migrated over the M-lineage-derived v17/18 body wall muscles and approached the somatic gonad, *egl-17::SL2::mNG::PH* was upregulated in dorsal uterine cells and P6.p but downregulated in other ventral midline cells (Fig. 2J). However, strong expression was maintained in V5-derived neurons (Fig. 2K). SMs crawled forwards using a broad protrusion at the leading edge and short filopodia as they migrated from their position over the v17/18 body wall muscles towards the *FGF/egl-17-* expressing dorsal uterine cell (Movie 1; Fig. S5). As SMs migrated over the somatic gonad and centered over the *FGF/egl-17-*expressing uterine progenitors (Fig. S6), *egl-17::SL2::mNG::PH* remained highly expressed in the dorsal uterine and P6.p cells with lower expression in other uterine cells, P5.p, P7.p, and other ventral midline cells (Fig. 2L, M). We did not observe cytoneme-like protrusions from *egl-17::mNG::PH-*expressing cells, although SMs were in close proximity to *FGF/egl-17-*expressing midline cells and neurons throughout much of their migration.

**Figure 2.**
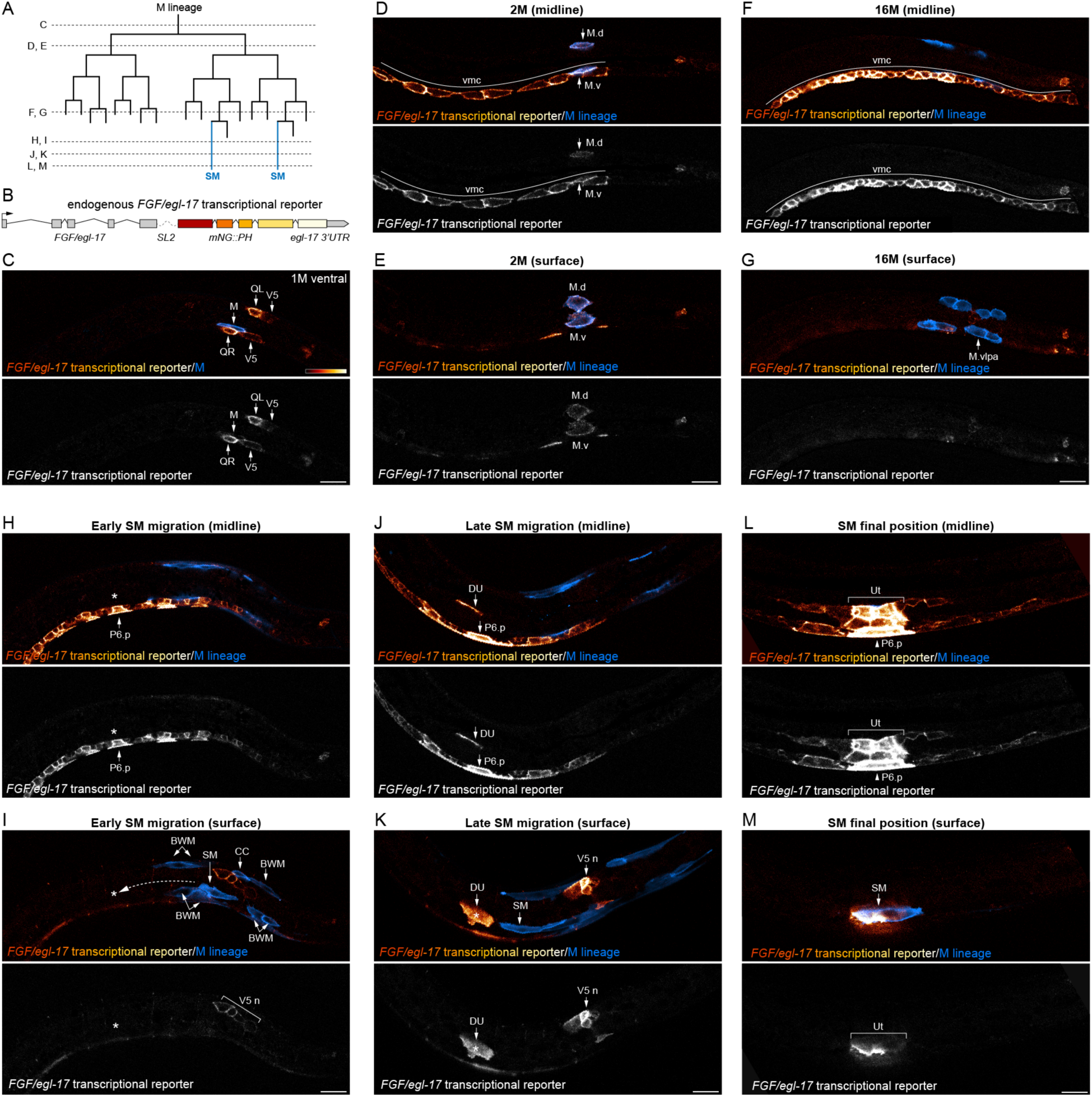
*In vivo* imaging of an endogenous *FGF/egl-17* transcriptional reporter during M lineage development reveals dynamic expression and multiple FGF/EGL-17 sources. (**A**) Schematic of M lineage divisions. Dashed lines indicate the developmental time points for images in C-M. (**B**) Design of the bicistronic, endogenous *FGF/egl-17* transcriptional reporter expressing *egl-17* and the plasma membrane marker mNG::PH under native regulatory control. (**C**) Ventral view of an early L1 stage animal showing *FGF/egl-17* expression in Q neuroblasts along with weak expression in V5 seam cells, a rectal cell, and M. (**D**, **E**) Lateral midline (**D**) and surface (**E**) views of a later L1 stage animal showing *FGF/egl-17* expression in ventral midline blast cells and the M lineage. (**F**, **G**) Midline (**F**) and surface (**G**) views at the 16M stage showing strong *FGF/egl-17* expression in a continuous row of ventral midline cells. The M.vlpa and M.vrpa cells divide to generate one SM and one additional body wall muscle on each side of the animal. (**H**, **I**) Midline (**H**) and surface (**I**) views showing *FGF/egl-17* expression in ventral midline cells including neurons and the presumptive VPCs during early SM migration. Asterisks denote the SM migration endpoint, and dashed arrow indicates the approximate extent of the FGF/EGL-17-dependent migration phase. Note that *FGF/egl-17* is also expressed in V5-derived neuroblasts dorsal to the position of the SM. (**J**, **K**) Midline (**J**) and surface (**K**) views showing *FGF/egl-17* expression in the dorsal uterine cell, VPCs, and V5-derived neurons during a late stage of SM migration. We did not observe cytonemes or other forms of direct contact between the dorsal uterine or P6.p cells and SMs during this FGF-dependent stage of migration. See Movie 1 and Fig. S5 for time-lapse imaging at this stage. (**L**, **M**) Midline and surface views showing strong *FGF/egl-17* expression in dorsal uterine cells and P6.p at the end of SM migration, along with weaker expression in other VPCs and somatic gonad cells. Animal in C is shown in ventral view with anterior to left. Animals in D-M are shown in lateral views and oriented with anterior to left and dorsal to top. Autofluorescent gut granules were removed from images by subtracting a 445ex/642-80em background channel. Abbreviations: BWM, body wall muscle; CC, coelomocyte; DU, dorsal uterine cell; SM, sex myoblast; Ut, uterine cells; V5n, V5-derived neuroblasts or neurons; vmc, ventral midline cells. Scale bars = 10 µm.

### FGF/EGL-17 orients migrating SMs

While *FGF/egl-17* expression at the end of SM migration resembled published transgenes, the endogenous *FGF/egl-17* transcriptional reporter revealed a more complex and dynamic expression pattern during the early larval development and SM migration. In particular, the finding that *FGF/egl-17* was expressed in the early M lineage and then in ventral midline cells along the path taken by migrating SMs raised the possibility that *FGF/egl-17* may act permissively in at least some aspects of SM migration rather than as an instructive signal. Given the available tools, earlier studies concluded that FGF/EGL-17 acts as a chemoattractant based on transgene expression patterns (Branda and Stern, 2000, Burdine et al., 1998) combined with the ability of SMs to follow the *FGF/egl-*17-expressing 1° VPC cells (Burdine et al., 1998) or the somatic gonad (Thomas et al., 1990) when they were misplaced. *FGF/egl-17* is required for the ability of 1° VPCs to mediate SM positioning in the absence of the somatic gonad (Burdine et al., 1998), but the use of genetic backgrounds including an overexpressed *Ras*/*let-60(G13E)* gain of function allele and/or laser ablation of the somatic gonad or VPCs complicates our understanding of the normal role for FGF/EGL-17 in guiding SM migration. Our results showing that *FGF/egl-17* is strongly expressed in ventral midline cells before VPC induction and in V5-derived neuroblasts at the end of SM migration raise the possibility that FGF/EGL-17 is required permissively for SMs to migrate anteriorly and SM positioning depends on a complementary “stop here” signal in wild-type animals. To distinguish between instructive and permissive roles for FGF in orienting migrating SMs, we used single copy transgenes to drive *FGF/egl-17* expression cell autonomously in M lineage cells, at uniform levels along the anteroposterior axis, or in tail cells in *FGF/egl-17(Δ)* (Fig. 3) and wild-type (Fig. S7) animals.

**Figure 3.**
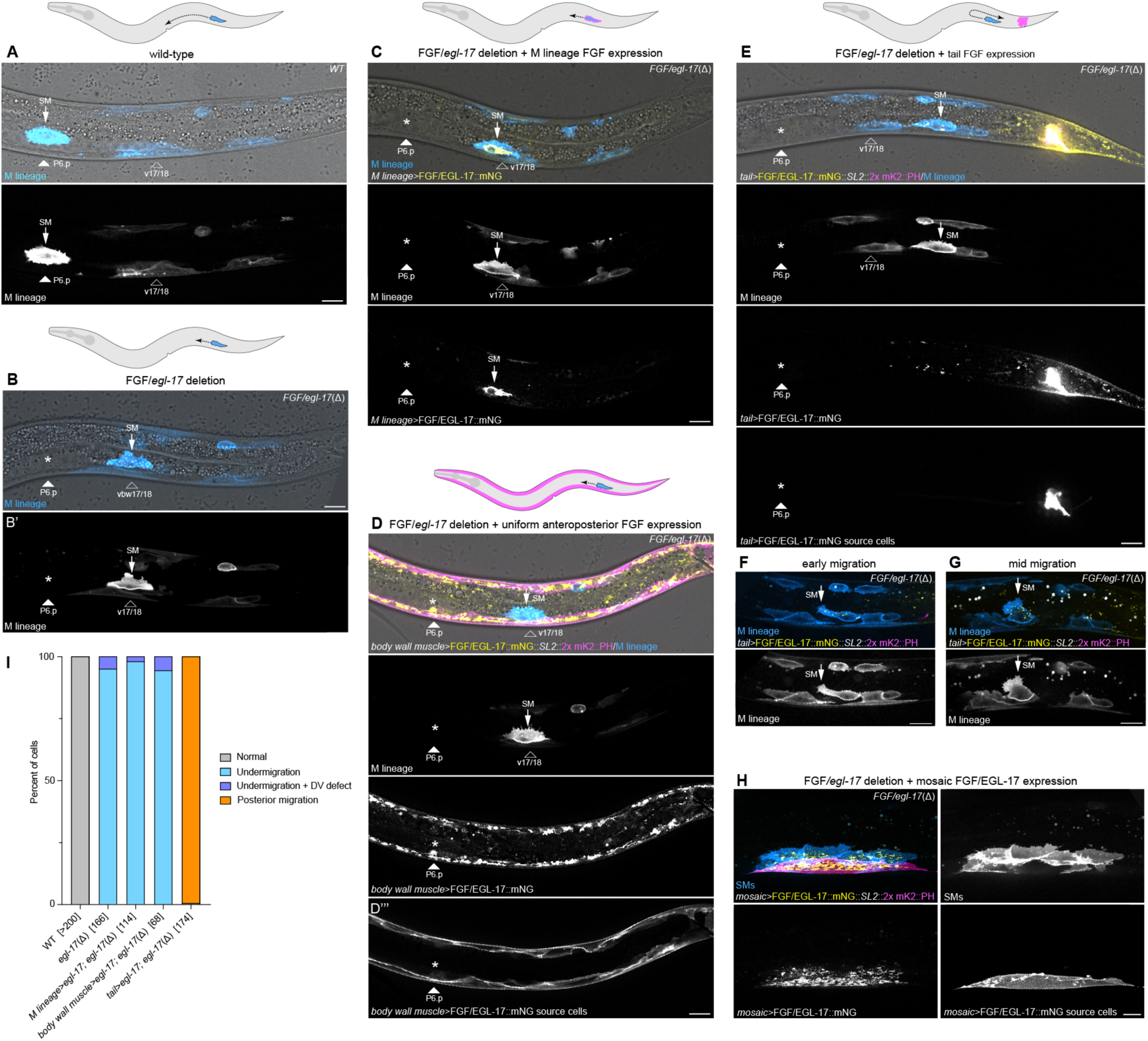
FGF/EGL-17 acts as an instructive signal to orient migrating SMs. (A) Final SM position over the somatic gonad center and P6.p in a wild-type animal. (B) Final SM positioning in a representative *FGF/egl-17(Δ)* animal showing migration arrest in the vicinity of the v17/18 body wall muscles. Asterisk marks the normal migration endpoint. (C) Expressing FGF/EGL-17::mNG in M lineage cells using the *hlh-8* promoter does not rescue SM migration in a *FGF/egl-17(Δ)* background. (D) uniform expression of FGF/EGL-17::mNG along the anteroposterior axis using the *myo-3* promoter does not rescue SM migration in a *FGF/egl-17(Δ)* background. (E) Expressing FGF/EGL-17::mNG in tail cells using a fragment of the *egl-20* promoter in a *FGF/egl-17(Δ)* animal leads to more posterior SM positioning. (F-G) SMs initially migrate anteriorly in *Pegl-20 ^-1261-610^ >FGF/egl-17::mNG; FGF/egl-17(Δ)* animals (F) but fail to maintain their polarity (G) before migrating towards the posterior. The tails cells expressing FGF/EGL-17::mNG are located out of the field of view to the right. (H) Isolated body wall muscles cells expressing FGF/EGL-17::mNG in mosaic animals can position SMs in the *FGF/egl-17(Δ)* background. (I) Summary of SM positioning phenotypes. Numbers in brackets indicate the number of cells scored for each genotype with two cells per animal (left and right). See Table S1 for source data. Asterisks denote the normal SM migration endpoint in images of cells with migration defects. 2x mKate2::PH marks membranes of the FGF-expressing cells in D-H. All images oriented with anterior to left and dorsal to top. Abbreviations: SM, sex myoblast; vbw, ventral body wall muscle. Autofluorescent gut granules were removed in B-E by subtracting a 445ex/642-80em background channel. Scale bars = 10 µm.

We reasoned that if FGF acts permissively in SM migration, then expressing *FGF/egl-17* cell-autonomously or at even levels throughout the body should rescue SM migration in *FGF/egl-17(Δ)* animals. We first used the *hlh-8* promoter to drive *FGF/egl-17::mNG* expression in M lineage cells including the SMs (Fig. 3C). *Phlh-8>FGF/egl-17::mNG* failed to rescue SM migration (n=114/114) in *FGF/egl-17(Δ)* animals (Fig. 3C, I), indicating that cell-autonomous FGF expression is not sufficient for SM migration. As a complementary test for permissive signaling, we used the *myo-3* promoter to express *FGF/egl-17::mNG* at uniform levels along the anteroposterior body axis. *Pmyo-3>FGF/egl-17::mNG* expression similarly failed to rescue SM migration (n=68/68) in *FGF/egl-17(Δ)* animals (Fig. 3D, I). To test the extent to which FGF acts as a chemoattractant for migrating SMs, we then used a fragment of the *egl-20* promoter to drive *FGF/egl-17::mNG* in tail cells in an attempt to reorient migrating SMs towards the posterior. In *FGF/egl-17(Δ)* animals, SMs typically arrest their migration over the v17/18 body wall muscle cells (Fig. 3B). Expressing *FGF/egl-17::mNG* in the tail led to posteriorly displaced SMs (n=173/174) that were located near the most posterior M-derived body wall muscles (Fig 3E, I). Because this position is near the birthplace of the SMs (see Fig. 1), we used live imaging at earlier timepoints to assess if the initial FGF-independent phase of migration failed. Intriguingly, we observed that initial SM migration occurred normally, and cells generated oriented protrusions and migrated anteriorly until reaching v17/18 (Fig. 3F). However, SMs subsequently failed to maintain their polarity and produced misoriented protrusions (Fig. 3G) before migrating posteriorly. Taken together, these results demonstrate that FGF acts instructively to orient SMs during the FGF-dependent phase of migration and that SMs are insensitive to an ectopic FGF source until the point when their migration normally requires FGF.

To test the ability of ectopic FGF sources to interfere with normal SM migration, we performed complementary misexpression experiments in a wild-type background. Cell-autonomous FGF/EGL-17 expression strongly disrupted SM migration (n=75/80), leading to undermigration reminiscent of FGF loss of function (Fig. S7). Uniformly expressing FGF/EGL-17 in body wall muscle cells led to similar undermigration defects in the vast majority (n=102/104) of cells (Fig. S7). High levels of FGF/EGL-17 expression in the tail disrupted SM migration in most animals (n=35/58), leading to cells that migrated towards the anterior but did not complete their migration (n=9/58) and cells that reoriented and migrated towards the posterior (n=26/58) (Fig. S7). Together, these findings confirm that FGF/EGL-17 acts instructively to guide migrating SMs and suggest that these cells are capable of considering the relative strengths of opposing FGF/EGL-17 sources.

### *FGF/egl-17(Δ)* somatic gonad cells can mediate precise SM positioning

To further explore the ability of FGF/EGL-17 to attract migrating SMs, we used mosaic *Pmyo-3>FGF/egl-17::mNG* expression to generate animals with ectopic point sources of FGF/EGL-17 in varying positions in the *FGF/egl-17(Δ)* background. In many cases, SMs centered over ectopic FGF/EGL-17-producing body wall muscles when these were located in the posterior half of the animal (Fig. 3H). However, in rare cases where mosaic FGF/EGL-17-secreting cells were in a similar anteroposterior position as the somatic gonad (n=4), we observed that SMs extended towards, or centered over, the somatic gonad rather than the ectopic FGF-expressing body wall muscle cell(s) (Fig. 4A). To test the extent to which somatic gonad cells are capable of capturing SMs migrating towards an ectopic FGF source without the caveats associated with mosaic overexpression, we expressed *FGF/egl-17* in the pharynx in an *FGF/egl-17(Δ)* background using a single copy transgene driven by a *myo-2* promoter (Fig. 4B). Anterior *FGF/egl-17* misexpression led to complex SM migration phenotypes including overmigration (42% of cells), normal positioning at the somatic gonad center (39%), and undermigration (19%) (Fig. 4B-E). SMs that stopped at their normal position maintained that position throughout proliferation and differentiation (Fig. S8). These results suggest that *FGF/egl-17(Δ)* somatic gonad cells provide a yet to be identified, short-range cue that can override conflicting positional information from an ectopic FGF source.

**Figure 4.**
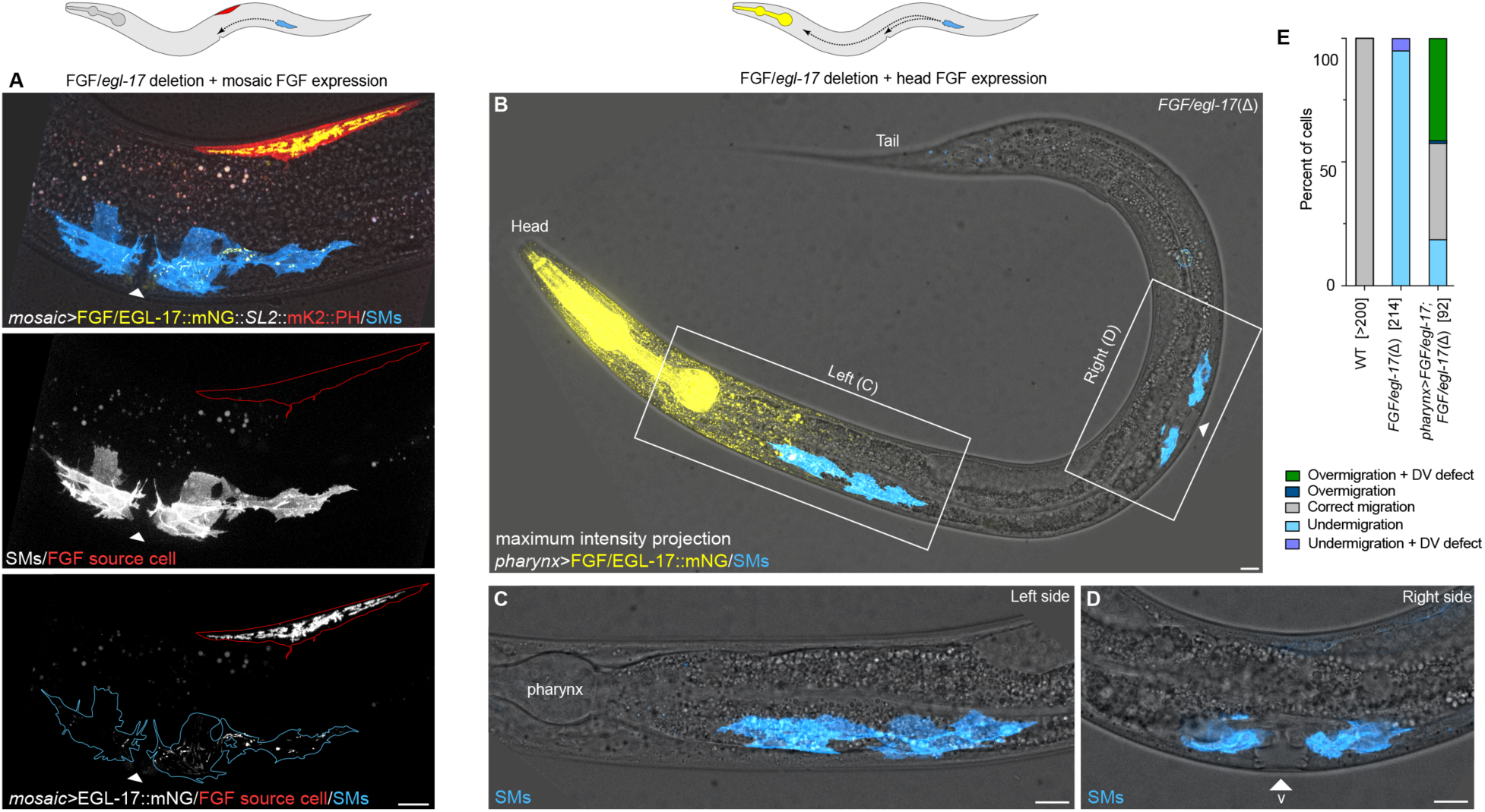
*FGF/egl-17(Δ)* gonad cells can capture SMs migrating towards a misplaced FGF/EGL-17 source. (A) Proper SM descendant positioning in a rare animal with mosaic *Pmyo-3>FGF/EGL-17::mNG* expression in a dorsal body wall muscle near the somatic gonad center. The SM descendants are located in their normal position on the ventral side flanking the somatic gonad center rather than near the FGF/EGL-17::mNG-expressing cell, marked by co-expressed 2x mKate2::PH. Note aberrant morphogenesis with a single SM daughter cell slightly displaced towards the posterior. (B) Misexpressing FGF/EGL-17::mNG in the head in an *FGF/egl-17(Δ)* background attracts migrating SMs, but *FGF/egl-17(Δ)* somatic gonad cells can also capture migrating SMs and mediate normal positioning. B shows a maximum intensity projection through an entire *Pmyo-2>FGF/EGL-17::mNG*; *FGF/egl-17(Δ)* animal where the left SM overmigrated, and the right SM is normally positioned. White triangle marks the normal SM migration endpoint. (C, D). Higher magnification views of overmigrated (C) and normally positioned (D) SM descendants in the same animal. White boxes in B indicate the regions shown in C and D. (E) Summary of SM positioning phenotypes in wild-type, *FGF/egl-17(Δ)*, and *Pmyo-2>FGF/EGL-17::mNG*; *FGF/egl-17(Δ)* animals. Numbers in brackets indicate the number of cells scored for each genotype with two cells per animal (left and right). See Table S1 for source data. All images are oriented with anterior to left and dorsal to top. Autofluorescent gut granules were removed in B-D by subtracting a 445ex/642-80em background channel. Scale bars = 10 µm.

### FGF/EGL-17 does not form a visible protein gradient *in vivo*

Understanding how secreted signaling proteins disperse requires the ability to visualize endogenous proteins in their native tissue context to determine how far ligands spread and assess the existence of signaling protein gradients. While FGF gradients are hypothesized to regulate many developmental processes, endogenous FGF protein gradients have only been directly observed in a handful of cases. To visualize FGF/EGL-17 protein in living animals, we used Cas9-triggered homologous recombination to endogenously tag EGL-17 at its C-terminus with mNG::3xFlag (hereafter EGL-17::mNG). EGL-17::mNG animals were phenotypically indistinguishable from wild-type, and FGF/EGL-17::mNG was visible at low levels throughout SM migration using spinning disk live imaging. We primarily observed FGF/EGL-17::mNG protein localized to cell types that expressed the *FGF/egl-17::SL2::mNG::PH* transcriptional reporter (see Fig. 2) and in M lineage cells, along with isolated punctae associated with other cell types (Fig. 5A-F). During the FGF-independent phase of SM migration, we observed FGF/EGL-17::mNG internalized within SMs and other M lineage cells (Fig. 5B). As SMs continued to migrate, we observed increasing accumulation of intracellular FGF/EGL-17::mNG in SMs with little visible fluorescence in additional cell types (Fig. 5C-F). Unexpectedly, FGF/EGL-17::mNG protein did not form a visible, extracellular concentration gradient at any stage. Because FGF/EGL-17::mNG fluorescence was only slightly higher than background, we used autofluorescence subtraction (Fig. S3) to visualize background-free localization of FGF/EGL-17::mNG at the onset of the FGF-dependent stage of SM migration. Maximum intensity projections of background-subtracted images showed FGF/EGL-17::mNG localization within the dorsal uterine cell, P6.p, ventral midline cells, and V5-derived neuroblasts (Fig. 5D) consistent with our *egl-17::SL2::mNG::PH* transcriptional reporter. Aside from *FGF/egl-17-* expressing cells, we observed bright punctae within migrating SMs and isolated punctae associated with other cell types (Fig. 5D). Even with autofluorescence subtraction, we did not observe an FGF/EGL-17::mNG gradient or protein associated with basement membranes or extracellular space. Although the absence of a clear FGF/EGL-17 protein gradient was unanticipated, this result in fact matches expectations for a freely diffusing protein in the absence of a uniformly expressed extracellular binding partner.

**Figure 5.**
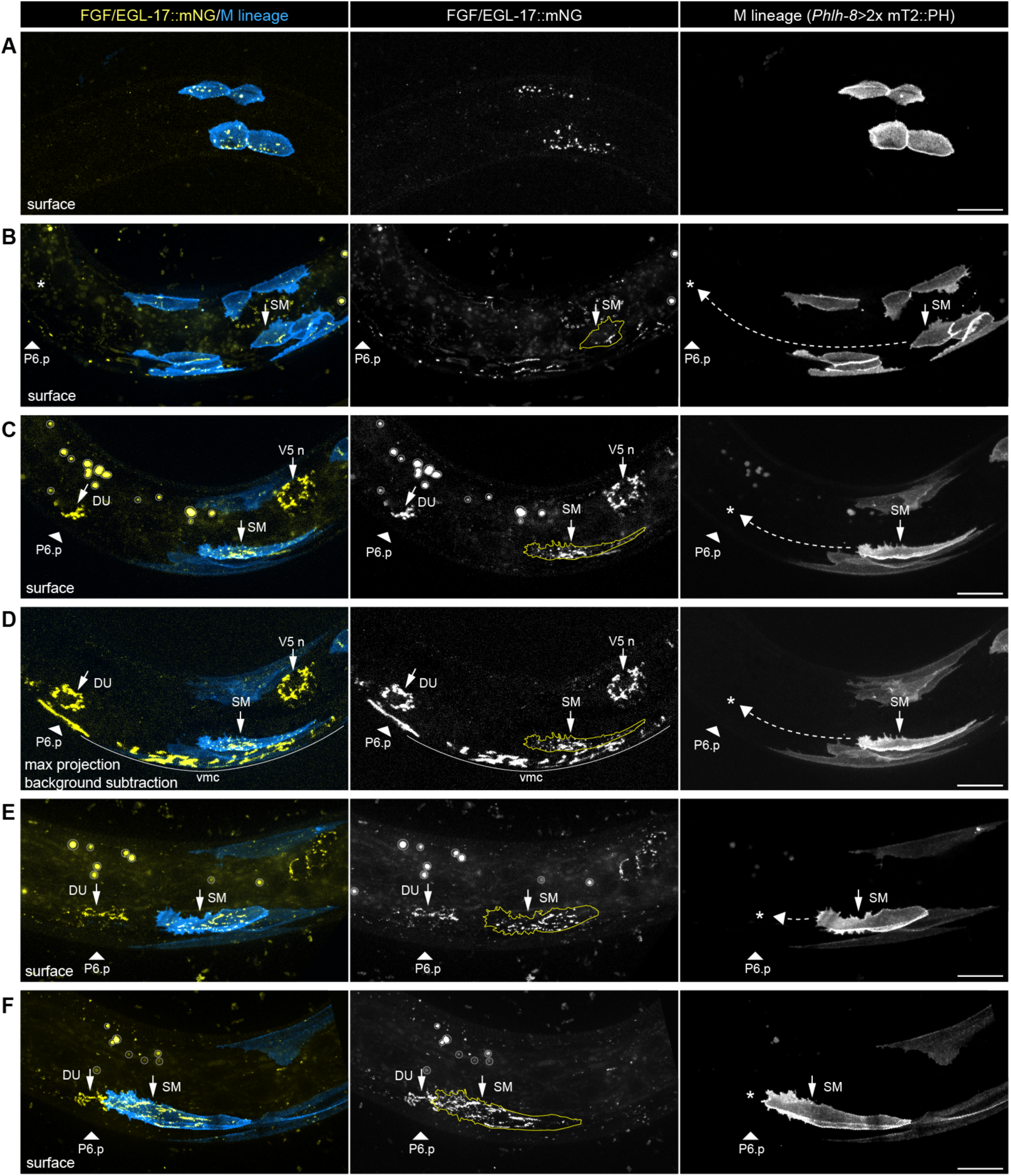
*in vivo* visualization of endogenously tagged FGF/EGL-17::mNG during SM migration. (**A**) Surface view showing FGF/EGL-17::mNG localizes to intracellular punctae in M lineage cells prior to SM birth. (**B**) Surface view showing that shortly after birth of the SMs, FGF/EGL-17::mNG localizes primarily to M lineage cells with sparse signal associated with other cell types. Dashed arrow indicates the path of SM migration, and asterisk indicates the endpoint. (**C**) Surface view showing FGF/EGL-17::mNG localization in the plane of SM migration at the beginning of the FGF-dependent phase of SM migration. FGF/EGL-17::mNG is visible as small punctae within the migrating SM and within the FGF/EGL-17-secreting dorsal uterine cell and V5-derived neuroblasts. (**D**) Maximum intensity projection of background-subtracted images showing FGF/EGL-17::mNG localization in the same animal as in C. Strong, intracellular FGF/EGL-17::mNG signal is visible in the SM along with the *FGF/egl-17*-expressing dorsal uterine cell, P6.p and other ventral midline cells, and V5-derived neuroblasts. Note the absence of an observable protein gradient between the migrating SM and its destination and only sparse FGF/EGL-17::mNG localization to other cell types. (**E**) Surface view showing FGF/EGL-17::mNG localization in the plane of SM migration prior to contact between the SM and dorsal uterine cell. (**F**) Surface view showing FGF/EGL-17::mNG localization in the plane of SM migration after contact between the SM and dorsal uterine cell. FGF/EGL-17::mNG is visible within the SM and in isolated punctae associated with other cell types. All images are oriented with anterior to left and dorsal to top. Asterisks denote the presumptive endpoint of SM migration in single channel images of *Phlh-8>2x mTurquoise2::PH.* Autofluorescent gut granules appear as circular artifacts in gut cells that are larger than FGF/EGL-17::mNG punctae and are outlined in gray in C, E, and F. Abbreviations: DU, dorsal uterine cell; SM, sex myoblast; V5 n, V5-derived neuroblasts or neurons; vmc, ventral midline cells. Scale bars = 10 µm.

### FGF/EGL-17 diffusion is required for SM migration

To elucidate potential requirements for FGF/EGL-17 diffusion in SM migration, we sought to prevent extracellular dispersal of FGF/EGL-17 without disrupting contact-dependent signaling. We first engineered an endogenously membrane-anchored FGF/EGL-17 by inserting mNG fused to a NLG-1 transmembrane domain (Wang et al., 2014, Teichmann and Shen, 2011) at the 3’ end of *FGF/egl-17* using Cas9-triggered homologous recombination in a wild-type background (Fig. 6A, B). *FGF/egl-17::mNG::nlg-1^TM^* animals displayed fully penetrant defects in SM migration (n=162/162; Fig. 6B, D), demonstrating that endogenously membrane-anchored FGF/EGL-17 is incapable of supporting normal SM migration. In addition to undermigration, we observed dorsoventral migration defects in a small number of animals along with a single case of overmigration (Fig. 6D). To confirm that membrane-anchored FGF/EGL-17::mNG::NLG-1^TM^ remained capable of contact-dependent signaling, we used the *myo-3* promoter to express *FGF/egl-17::mNG::nlg-1^TM^* in body wall muscles, which directly contact SMs. *Pmyo-3>FGF/egl-17::mNG::nlg-1^TM^* animals exhibited SM migration defects (n=98/98; Fig. S9) similar to the phenotype caused by un-tethered FGF/EGL-17 misexpression in body wall muscles (Fig. S7; Table S1), thereby validating that FGF/EGL-17::mNG::NLG-1^TM^ was competent to activate signaling at cell contacts.

**Figure 6.**
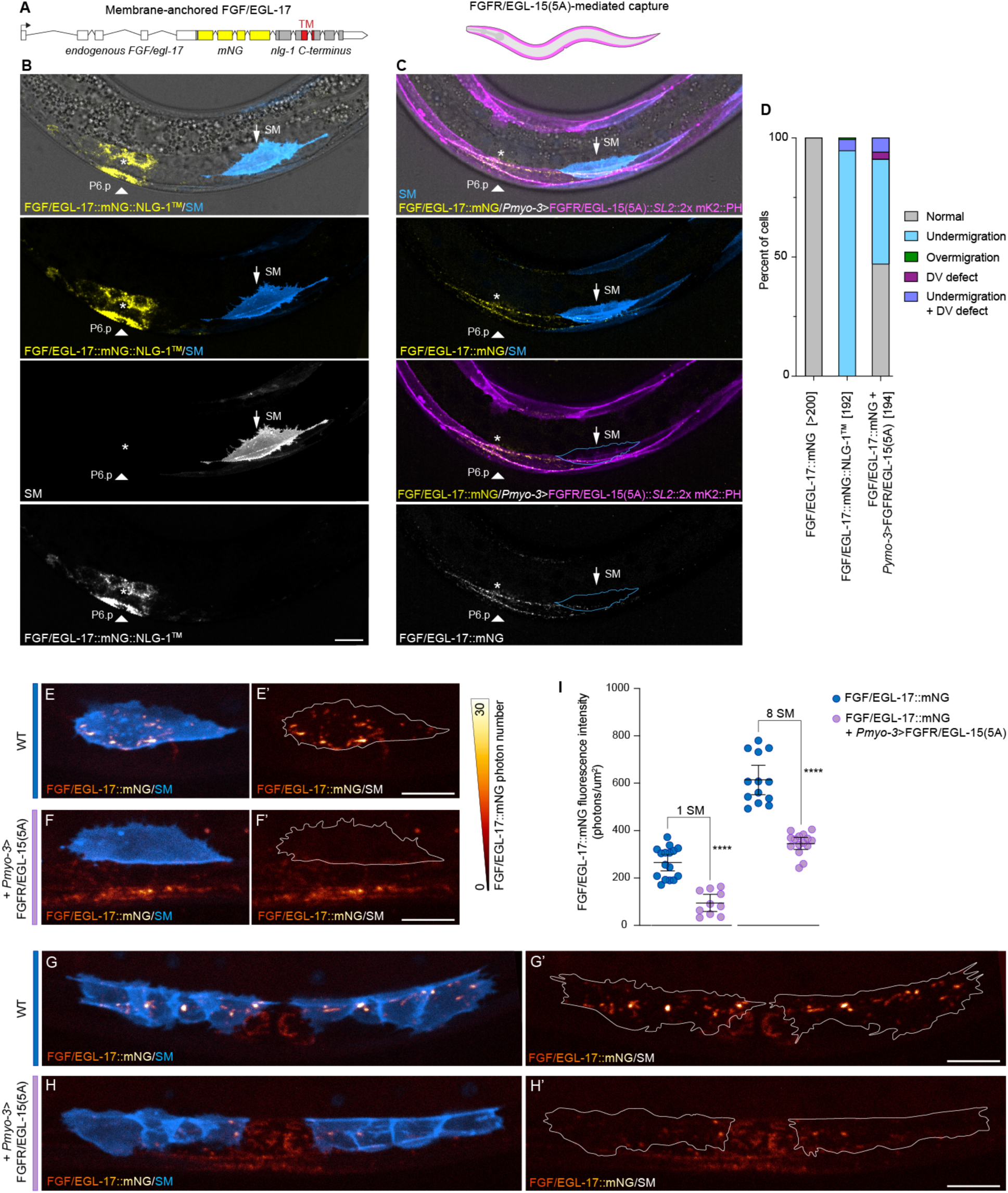
FGF/EGL-17 diffusion is required for SM migration. (**A**) Schematic of the endogenously engineered membrane-tethered FGF/EGL-17 allele. Endogenous *egl-17* coding and 3’UTR sequences are indicated in white, *mNG* is shown in yellow, and the *nlg-1* C-terminus is shown in gray with the transmembrane domain indicated in red. (**B**) FGF/EGL-17::mNG::NLG-1^TM^ is not capable of supporting SM migration. White triangle indicates P6.p and asterisk indicates the normal endpoint of SM migration. (**C**) Expressing FGFR/EGL-15(5A) in body wall muscles sequesters endogenous FGF/EGL-17::mNG and disrupts SM migration. Note that endogenous FGF/EGL-17::mNG localizes to body wall muscles near *FGF/egl-17*-expressing cells in *Pmyo-3>FGFR/EGL-15(5A)::SL2::2x mKate2::PH; FGF/egl-17::mNG* animals but not the SM. White triangle indicates P6.p and asterisk indicates the normal endpoint of SM migration. (**D**) Summary of SM positioning phenotypes in *FGF/egl-17::mNG, FGF/egl-17::mNG::nlg-1^TM^*, and *Pmyo-3>FGFR/EGL-15(5A)::SL2::2x mKate2::PH* animals. Numbers in brackets indicate the number of cells scored for each genotype with two cells per animal (left and right). See Table S1 for source data. (**E**, **F**) Comparison of FGF/EGL-17::mNG localization in SMs at the end of migration in an *FGF/egl-17::mNG* animal (**E**) and a *Pmyo-3>FGFR/EGL-15(5A)::SL2::2x mKate2::PH; FGF/egl-17::mNG* with normal SM migration (**F**). FGF/EGL-17 captured by body wall muscles is visible ventral to the SM (**G**, **H**) Comparison of FGF/EGL-17::mNG localization in SMs at the 8 SM stage in an *FGF/egl-17::mNG* animal (**G**) and a *Pmyo-3>FGFR/EGL-15(5A)::SL2::2x mKate2::PH; FGF/egl-17::mNG* animal with normal SM migration (**H**). (**I**). Quantification of FGF/EGL-17::mNG levels in SMs in control and *Pmyo-3>FGFR/EGL-15(5A)::SL2::2x mKate2::PH* animals with normal migration. Expressing FGFR/EGL-15(5A) in body wall muscles significantly reduces FGF/EGL-17::mNG even in SMs that correctly migrated (Mann-Whitney test, P<0.0001 at both time points). All images are oriented with anterior to left and dorsal to top. Images in E-H and images used for quantification in I were acquired using a Hamamatsu ORCA Quest quantitative CMOS camera in its photon number resolving mode. Autofluorescent gut granules were removed in B and C by subtracting a 445ex/642-80em background channel. Scale bars = 10 µm.

As a parallel approach to test the extent to which FGF/EGL-17 diffusion is required for SM migration, we sought to limit extracellular FGF/EGL-17 dispersal without functionally modifying the protein itself. To do so, we first attempted to sequester free, extracellular FGF/EGL-17 using an anti-GFP nanobody-based approach known as Morphotrap (Harmansa et al., 2015) similar to that used to test roles for FGF8a diffusion in zebrafish embryos (Harish et al., 2023). We tagged endogenous FGF/EGL-17 with the GFP-derivative YPET and crossed FGF/EGL-17::YPET animals to a Morphotrap strain that was previously used to demonstrate a requirement for WNT/EGL-20::YPET diffusion in long-range signaling (Pani and Goldstein, 2018). However, Morphotrap appeared unable to capture FGF/EGL-17::YPET, which did not localize to Morphotrap-expressing body wall muscle cells despite their proximity to both the FGF/EGL-17::YPET-expressing cells and SMs (Fig. S10). As expected given its failure to sequester FGF/EGL-17::YPET, *Pmyo-3>*Morphotrap had no effect on SM migration (Fig. S10, Table S1; n=76/76).

As an alternative approach to capture the diffusing fraction of FGF/EGL-17 protein, we used a single-copy transgene to express *FGFR/egl-15(5a)* along with a *2x mKate2::PH* membrane marker in body wall muscles to provide a receptor-mediated sink for extracellularly dispersing FGF/EGL-17::mNG (Fig. 6C). We did not observe defects in body wall muscle differentiation or anatomy visualized by the 2x mKate2::PH membrane marker in this strain. While endogenously tagged FGF/EGL-17::mNG does not localize to body wall muscles during SM migration in wild-type animals (see Fig. 5C-F), we observed conspicuous FGF/EGL-17::mNG localization to body wall muscles near *FGF/egl-17-*expressing cells in *FGF/egl-17::mNG; Pmyo-3>FGFR/egl-15(5a)::SL2::2x mKate2::PH* animals (Fig. 6C; Fig. S11). Consistent with the ability of this transgene to capture FGF/EGL-17::mNG, we observed migration defects in the majority of SMs (n=101/194; Fig. 6B, C). Like *FGF/egl-17* deletion phenotypes, we primarily observed undermigration along with rarer dorsoventral defects (Fig. 6D; Fig. S11; Table S1). Because the *FGFR/egl-15(a)*-misexpressing body wall muscles are not directly juxtaposed between the SMs and *FGF/egl-17*-expressing cells, we hypothesized that *FGFR/egl-15(5a)* misexpression interfered with SM migration by sequestering FGF/EGL-17 from a common pool of extracellularly diffusing protein, thereby reducing the amount of extracellular FGF available to SMs below the quantity required for migration. To assess this model, we used spinning disk live imaging with a photon number-resolving qCMOS camera to quantify the amount of FGF/EGL-17::mNG internalized by SMs in wild-type and *Pmyo-3>FGFR/egl-15(5a)* animals (Fig 6E-I). To avoid potentially confounding effects of cell position, we only utilized cells that migrated to their normal position for comparisons. Even when SMs migrated to their normal positions, we found that *Pmyo-3>FGFR/egl-15(5a)* significantly reduced the amount of FGF/EGL-17::mNG internalized by SMs at both the 1 SM (Fig. 6E, F, I) and 8 SM stages (Fig. 6G-I) (Mann-Whitney test P<0.0001 at both stages). This result confirmed that SMs acquire FGF/EGL-17 from a general pool of extracellularly mobile protein even though SMs at the 1 SM stage directly overlie *FGF/egl-17*-expressing cells (see Fig. S6). This finding further highlights that only vanishingly small quantities of ligand are required for FGF-dependent SM migration.

## Discussion

*In vivo* visualization of endogenous FGF/EGL-17 and experimental manipulations of extracellular dispersal, along with visualization of signaling and responding cell dynamics, provide direct evidence that FGF/EGL-17 is natively diffusible and that free, extracellular FGF dispersal is required for *C. elegans* muscle progenitor migration. While the FGF signaling pathway is evolutionarily conserved, analyses of endogenously tagged FGF proteins have shown disparate mechanisms for FGF movement between cells and gradient formation in vertebrates (Harish et al., 2023, Dubrulle and Pourquie, 2004, Yu et al., 2009) and *Drosophila* (Du et al., 2018, Du et al., 2022, Stepanik et al., 2020). In developing zebrafish, FGF8a diffuses extracellularly to form long-range protein gradients (Harish et al., 2023), and multiple studies have observed FGF diffusion in vertebrate models using overexpression-based approaches (Toyoda et al., 2010, Yu et al., 2009, Duchesne et al., 2012), with the associated caveats. However, the *Drosophila* FGFs characterized to date do not freely spread between cells by diffusion but instead signal at cell contacts (Du et al., 2022, Du et al., 2018, Stepanik et al., 2020) and form gradients through a cytoneme-based mechanism in cases where gradients exist (Du et al., 2018). Our results demonstrating that *C. elegans* FGF/EGL-17 moves between cells by diffusion, and that diffusion is required for at least one of its key functions, suggests that some invertebrate FGFs are endogenously diffusible and that this dispersal mechanism is a common feature of FGF proteins across phyla. Interestingly, the *Drosophila* FGFs that signal at cell contacts include extended C-terminal domains with a transmembrane domain (Stepanik et al., 2020) or GPI anchor (Du et al., 2022) that are absent in *C. elegans* FGF/EGL-17 and vertebrate FGF8/17/18 family proteins. However, the *C. elegans* FGF9/16/20 homolog LET-756 also includes a large C-terminal domain, and studies investigating FGF/LET-756 signaling mechanisms may provide additional insights into the diversity of mechanisms used for FGF dispersal.

While our findings argue that FGF/EGL-17 signaling in SM migration relies on extracellular diffusion, they do not rule out roles for cytonemes in other aspects of SM development or contact-dependent FGF/EGL-17 signaling in differing contexts. Indeed, the observation that stationary SMs extend numerous filopodia after migration has ended and throughout proliferation and morphogenesis could be consistent with a role for cytonemes in communication between SMs and other cell types at later developmental stages. Modes of signaling protein dispersal may also vary depending on tissue architectures (Stapornwongkul and Vincent, 2021), which affect the abilities of signaling and responding cells to make direct contacts. During SM migration, somatic gonad cells are encased within a basement membrane that would interfere with cytoneme-based signaling between the FGF/EGL-17-expressing uterine cells and migrating SMs. However, the fact that FGF8a moves by extracellular diffusion in early zebrafish development (Harish et al., 2023) demonstrates diffusion can be the predominant mode for FGF dispersal even when there are not physical barriers to cell contacts.

*C. elegans* SMs are a long-standing paradigm to study directed cell migration (Chen and Stern, 1998, Sherwood and Plastino, 2018), but technical limitations prevented early studies (Burdine et al., 1998) from directly testing whether FGF/EGL-17 functions as a chemoattractant that guides migrating SMs towards the highest source of FGF or as a permissive signal that allows cells to migrate anteriorly until receiving a positioning cue from the somatic gonad or 1° VPCs. Our results using *FGF/egl-17* misexpression in an *FGF/egl-17(Δ)* background demonstrate that elements of both models are correct. FGF/EGL-17 is clearly not a permissive signal for SM migration as neither cell-autonomous nor spatially uniform *FGF/egl-17* expression are sufficient for SM migration in *FGF/egl-17(Δ)* animals, and both manipulations disrupt SM migration in a wild-type background. Misexpressing *FGF/egl-17* in tail cells reversed the direction of SM migration, demonstrating that extracellular FGF can orient migrating SMs towards a distant source, and FGF/EGL-17 functions as a bona fide chemoattractant for migrating SMs. However, sparse mosaic and pharyngeal *FGF/egl-17* expression also highlighted the ability of *FGF/egl-17(Δ)* somatic gonad cells to attract SMs over short distances, or to halt their migration when they are in close proximity. The molecules that mediate these interactions between SMs and the somatic gonad remain to be identified, and it is also not known if the ability of *FGF/egl-17(Δ)* gonad cells to precisely position migrating SMs depends indirectly on FGF/EGL-17, which was expressed in distant cells in these experiments. It is possible that the proteins that mediate short-range communication between the gonad and SMs are related to the gonad-dependent repulsive guidance signal that repels migrating SMs in the absence of FGF/EGL-17 or FGFR/EGL-15(5A) (Branda and Stern, 2000, Sherwood and Plastino, 2018), which also awaits discovery.

*C. elegans* has a single FGFR homolog, *egl-15*, whose extracellular domain is alternatively spliced to generate receptor isoforms with differential affinity for EGL-17 and the second *C. elegans* FGF ligand, LET-756 (Lo et al., 2008, Goodman et al., 2003). Our finding that expressing the FGFR/EGL-15(5A) isoform in body wall muscle leads to SM migration defects by sequestering extracellular FGF/EGL-17 suggests a novel biological rationale for multiple FGFR isoforms in *C. elegans*. FGF/LET-756 signaling in the hypodermis is required for larval viability, which is mediated by FGFR/EGL-15(5B). Although FGFR/EGL-15(5A) is capable of performing the essential functions of EGL-15(5B) (Lo et al., 2008), expressing levels of FGFR/EGL-15(5A) sufficient for FGFR/LET-756 signaling in the hypodermis could interfere with SM migration given that FGF/EGL-17 is produced at very low levels. In this case, alternative FGFR isoforms may have evolved in part to protect a low-expressed chemoattractant, FGF/EGL-17, from being functionally sequestered by FGFR-expressing cells that respond to the more abundant FGF/LET-756. It is possible that avoiding the ability to sequester related ligands and prevent their dispersal to target cells is an underappreciated function for receptor diversification.

FGFs can act as morphogens (Toyoda et al., 2010, Harish et al., 2023, Kengaku and Okamoto, 1995), and FGF protein concentration gradients are thought to be important for conveying positional information in developing animals. In classic models, spatial gradients of FGF protein are essential to activate distinct concentration (or fold change) -dependent responses in cells located at varying distances from an FGF source(Balasubramanian and Zhang, 2016). While we demonstrated that FGF/EGL-17 acts instructively to orient mesodermal progenitor migration in *C. elegans*, endogenous FGF/EGL-17 unexpectedly did not form a visible protein gradient *in vivo* during SM migration. Nonetheless, cell-autonomous, uniform, and posterior FGF/EGL-17 misexpression experiments demonstrated that a localized FGF/EGL-17 source is required for SM migration and migrating cells orient towards the strongest FGF source. Our discovery that endogenous FGF/EGL-17 is diffusible but does not form observable gradients *in vivo* may be explained by the fact that freely diffusible proteins require extracellular binding partners to form stable, long-range concentration gradients. Theoretical predictions and experimental tests using secreted GFP demonstrated that freely diffusible proteins do not form concentration gradients on their own *in vivo* (Stapornwongkul et al., 2020), and the length and shape of a secreted protein gradient depends on the distribution of binding partners and their affinities (Stapornwongkul et al., 2020, Stapornwongkul and Vincent, 2021, Muller et al., 2013). Interestingly, the *FGFR/egl-15(5a)* isoform that binds EGL-17 and is required for SM migration is expressed predominantly in the M lineage (Lo et al., 2008), suggesting that other nearby cell types may lack high-affinity binding partners for FGF/EGL-17. It is possible that the absence of an observable, endogenous FGF/EGL-17 gradient reflects a lack of extracellular binding partners expressed in cells outside the M lineage. Secreted protein gradients can also arise through binding to non-receptor binding partners with lower affinities (Stapornwongkul and Vincent, 2021, Stapornwongkul et al., 2020). In many contexts, FGFs associate with heparan sulfate proteoglycans (HSPGs) in the extracellular matrix (Duchesne et al., 2012, Muller et al., 2013, Balasubramanian and Zhang, 2016), but such interactions are not known for EGL-17, and viable *C. elegans* HSPG mutations have not been reported to affect SM migration. Collectively migrating cells can generate their own gradients in migrating tissues (Dona et al., 2013, Venkiteswaran et al., 2013), but such a mechanism has not been described for individually migrating cells *in vivo*. In the absence of an observable protein gradient, it is possible that intracellular signal transduction processes (Cai and Montell, 2014, Casaletto and McClatchey, 2012, Shellard and Mayor, 2020) amplify slight differences in extracellular FGF levels to orient migrating cells. The fact that substantially decreased levels of FGF/EGL-17 are still capable of mediating SM migration in some *Pmyo-3>FGFR/egl-15(5a)* animals highlights that extremely small quantities of FGF/EGL-17 are sufficient for this biological function. However, the FGF signal transduction mechanisms that govern directed cell migration in SMs downstream of FGF/EGL-17 remain uncertain due in part to pleiotropic functions of key signal transduction components (Sherwood and Plastino, 2018, Sundaram et al., 1996, Sundaram, 2013). Our work demonstrates that invertebrate FGFs may have been ancestrally diffusible and also lays the groundwork to investigate how migrating cells interpret multiple types of extracellular signals to navigate to precise destinations *in vivo*.

## Materials and Methods

### *C. elegans* maintenance

*Caenorhabditis elegans* animals were cultured on Nematode Growth Medium (NGM) plates, fed *E. coli* OP50, and maintained at 20°C for experiments. Only well-fed animals were used for live imaging and/or scoring SM positioning.

### Plasmid construction

Homologous repair templates for the FGF/EGL-17 locus were constructed by inserting homology arm PCR products into plasmids containing a self-excising selection cassette using New England Biolabs HiFi DNA assembly master mix as described in detail elsewhere (Dickinson et al., 2015). To construct the repair template used to delete the *FGF/egl-17* gene, we constructed an mNG^SEC^3xFlag repair template with homology arms that flanked the coding region to replace the entire *FGF/egl-17* coding sequence with mNG. To generate the repair template for the *FGF/egl-17* transcriptional reporter, we cloned *SL2::mNG::PH* in place of *mTurquoise2* in pDD315 and used the same homology arms as for endogenous protein tagging. A stop codon was added between the *FGF/egl-17* coding sequence and *SL2*. To generate the repair template for membrane-tethered FGF/EGL-17::mNG::NLG-1^TM^, we used the pDD268 backbone and the same homology arms as for FGF/EGL-17::mNG with the addition of a 2265 bp genomic fragment encoding the last 184 amino acids of NLG-1 between 3xFlag and the stop codon and 3’ homology arm. The NLG-1 C-terminus includes a single transmembrane domain and was previously used as a C-terminal membrane tether for functional, membrane-anchored Netrin/UNC-6 (Teichmann and Shen, 2011, Wang et al., 2014). Plasmid backbones for single copy insertions near the ttTi4348 and ttTi5605 sites were described previously (Pani and Goldstein, 2018). The plasmid backbone for single copy insertions at Chr IV:4,237,723 was constructed by cloning flanking genomic homology arms with the guide RNA target sequence deleted into pDD315 with the *mTurquoise2* and *3xHA* sequences removed. Coding sequences for transgenes were codon optimized using the *C. elegans* codon adapter tool (Redemann et al., 2011). Transgene promoters, fluorescent proteins, and the *SL2* sequence were amplified from genomic DNA or existing plasmids and inserted into linearized backbones using New England Biolabs HiFi DNA assembly. The *egl-20^(-1261−610)^* enhancer was cloned upstream of a *pes-10* minimal promoter. To construct plasmids for Cas9 + sgRNA expression, we cloned guide RNA sequences into the *Peft-3>Cas9 + sgRNA* expression vector pDD162 using site-directed mutagenesis. Guide RNA target sites used were (5’ – 3’): *egl-17* N-terminus, TCGACAACATCAGGGTGAGT; *egl-17* C-terminus, CACATGATAGTTTGTATCGT; Chr I transgenes, GAAATCGCCGACTTGCGAGG; Chr II transgenes, GATATCAGTCTGTTTCGTAA; Chr IV transgenes, ACTGTTGGATGCCTGTGTAG. Sequences for the *nlg-1* C-terminal region and transgene promoters used are provided in Supplemental File 1.

### Transgenesis and genome engineering

All strains generated here were made using Cas9-triggered homologous recombination with a self-excising selection cassette (SEC) (Dickinson et al., 2015). Single copy transgenes were inserted into chromosomal safe harbor locations near the ttTi4348 site on Chr I, the ttTi5605 site on Chr II, or a new site at Chr IV:4,237,723. FGF/EGL-17 was endogenously tagged at the C-terminus to generate mNG, YPET, and mNG::NLG-1^TM^ fusion alleles. The endogenous *FGF/egl-17* transcriptional reporter was engineered by inserting *SL2::mNG::PH* immediately after the stop codon using the same guide RNA. For transgenesis and endogenous tagging, we co-injected the homologous repair template, Cas9 + sgRNA plasmid, and extrachromosomal array markers into adult germlines as described previously (Dickinson et al., 2015, Dickinson et al., 2013). We handled animals with a fine paintbrush during the microinjection process to increase survival and throughput (Gibney et al., 2023). Candidate knock-in animals were selected using hygromycin B, a dominant roller phenotype, and the absence of fluorescent extrachromosomal array markers. To isolate homozygous lines, we picked single animals to new NGM plates without hygromycin B and screened for animals with 100% roller progeny. To excise the selectable marker cassette, we heat-shocked homozygous, young L1 larvae for four hours at 34°C to induce *heat shock>Cre* expression and picked non-roller offspring in the next generation. To generate strain APL670 for Morphotrap experiments, we crossed FGF/EGL-17::YPET animals with an existing *Pmyo-3>Morphotrap^SEC* strain (Pani and Goldstein, 2018) followed by heat shock to excise the SEC. To visualize ERK activity, we crossed ljfSi41; ljfSi42 animals to an existing *Phlh-8>ERK-nKTR* strain (de la Cova et al., 2017).

### Microscopy and image processing

Larval animals were immobilized using 0.1 mmol/L levamisole in M9 buffer (Chai et al., 2012) for standard live imaging or 0.1 µm polystyrene nanoparticles without anesthetic (Kim et al., 2013) for time-lapse imaging of cell migration. Animals were mounted between a high-precision cover glass and 3-5% (wt/vol) agarose pads. Animals were imaged within one hour of mounting, and images shown were representative of at least 10 animals. Numerous animals were mounted on each slide and selected for imaging based on developmental stage and orientation. Images were acquired using a Yokogawa CSU-X1 spinning disk confocal with a Hamamatsu ORCA Fusion BT sCMOS camera or a Yokogawa CSU-W1 SoRa spinning disk confocal with a Yokogawa Uniformizer and Hamamatsu ORCA Quest qCMOS camera. Both confocal units were mounted on Nikon Ti2 inverted microscope stands. Imaging was performed using 445 nm, 514 nm, 561 nm, or 594 nm lasers for excitation and 480/40, 545/40, 575LP, 610LP, or 642/80 emission filters depending on fluorescent protein. Imaging was performed using Nikon Plan Apo IR 60X/1.27 NA water immersion, Apo TIRF 60X/1.49 NA oil immersion, or Apo TIRF 100x/1.49 NA oil immersion objectives. Images were acquired using Nikon NIS Elements Advanced Research software (5.42.04 and 5.42.06) with camera exposure times and laser intensities adjusted based on experimental requirements and fluorescence intensities of the imaged proteins. For animals where the areas of interest were not located within a single field of view, we acquired tiled images using the large image acquisition mode with automated image stitching. Stitched images were not used for fluorescence intensity measurements. Images were deconvolved using Nikon NIS Elements software. Brightness, contrast, and colors were adjusted using Fiji (Schindelin et al., 2012), and figures were prepared using Adobe Illustrator 28.3.

*C. elegans* larvae and adults possess gut granules with broad-spectrum autofluorescence that can interfere with visualizing fluorescent proteins by live imaging. Gut granule autofluorescence is correlated across excitation and emission wavelengths (Rodrigues et al., 2022) and is prominent with all laser excitation wavelengths and emission filters used in this study. To better visualize cell architectures and weakly-expressed endogenous proteins, we subtracted gut granule autofluorescence in some images (see Fig. S3) using a method similar to SAIBR (Rodrigues et al., 2022). To subtract autofluorescence, we first acquired background images using 445 nm excitation paired with 642/80 or 575LP emission filters, which allowed us to spectrally separate broad-spectrum autofluorescence from the fluorescent proteins used here. We adjusted excitation power such that gut granules in the background channel were slightly more intense than gut granules in the fluorescent protein channels (445ex, 480/40em; 514ex, 545/40em; 561ex, 575LPem; 594ex, 642/80em) and then acquired paired background and fluorescent protein images at each z-position. We then subtracted the background channel from the fluorescent protein channel(s) using the “subtract” option in the Fiji Image Calculator (see Fig. S3). In cases where gut granules in the background channel were substantially brighter than in a fluorescent protein channel, we subtracted a constant from all pixels in the background channel to roughly equalize gut granule fluorescence intensity across channels before subtracting the background channel.

### Quantitative analyses

We scored L3 or L4 animals for SM migration defects using *Phlh-8*-driven transgenes to visualize SMs at developmental stages when they have normally completed their migration. At the 1 SM stage, we scored SMs as normally migrating when the single SM was centered over the uterine cells and P6.p. For animals imaged at later developmental stages, we scored SMs as normally migrating when the SM progeny were centered over the somatic gonad and symmetrically located flanking the P6.p descendants or the vulva. Cells that migrated anterior to P6.p were scored as overmigrating. Cells that were located posterior to P6.p but anterior to the most posterior M-derived body wall muscles were scored as undermigrating. Cells located over, or posterior to, the most posterior M-derived body wall muscles were scored as posteriorly migrating. To quantify FGF/EGL-17::mNG levels, we imaged animals using a Hamamatsu ORCA Quest qCMOS camera in its photon number resolving mode for absolute measurements of fluorescence intensity. To calculate FGF/EGL-17::mNG photon number, we drew a region of interest using the *Phlh-8>2x mTurquoise2::PH* membrane marker and measured FGF/EGL-17::mNG photon number in a z-projection of image planes encompassing the SM. Statistical significance of the difference in fluorescence intensity between wild-type and *Pmyo-3>FGFR/egl-15(5a)::SL2::2x mKate2::PH* backgrounds was assessed using a Mann-Whitney test. To quantify ERK-nKTR activity, we calculated the ratio of ERK-nKTR nuclear to cytoplasmic fluorescence intensity using co-expressed mCherry::H2B to visualize the nucleus (de la Cova et al., 2017). For ERK-nKTR measurements, we subtracted off-worm background in a nearby region from the raw pixel intensities. We assessed statistical significance using a Kolmogorov Smirnov test. All statistical tests were performed using Graphpad Prism 10.2.2 software. No animals or data points were excluded *post hoc*.

## Acknowledgements

Some strains were provided by the CGC, which is funded by the National Institutes of Health Office of Research Infrastructure Programs [P40 OD010440]. pRCA5 was a gift of Rebecca Adikes. This work was supported by the National Institutes of Health [Grant R35GM142880 to A.M.P. and Predoctoral Fellowship F31HD112152 to T.V.G.].

## Author Contributions

Conceptualization, TVG and AP; methodology, TVG and AP; formal analysis, TVG and AP; investigation, TVG and AP; writing – original draft, TVG and AP; writing – review and editing, AP; visualization, TVG and AP; funding acquisition, TVG and AP.

## Resource Availability

*C. elegans* strains generated in this study will be deposited at the *Caenorhabditis* Genetics Center (CGC) (https://cgc.umn.edu/) upon manuscript acceptance. Plasmids and other reagents will be shared on request.

## Supplemental Information

**Movie 1. Time-lapse imaging of SM migration in concert with an endogenous *FGF/egl-17* transcriptional reporter.** Spinning disk confocal time-lapse showing a portion of the FGF-dependent phase of SM migration. The SM and other M lineage cells are visualized in cyan. The *FGF/egl-17* transcriptional reported is depicted using the glow lookup table. Movie corresponds to Fig. S5. Animal is oriented with anterior to left and dorsal to top. Time elapsed is labeled in minutes. Scale bar = 10 µm. Movie at Figshare:https://doi.org/10.6084/m9.figshare.28611896.v1

