## Supplemental Information for "FGF diffusion is required for directed migration of postembryonic muscle progenitors in *C. elegans*"

**Figure S1. The gonadal basement membrane separates migrating SMs from somatic gonad cells.**

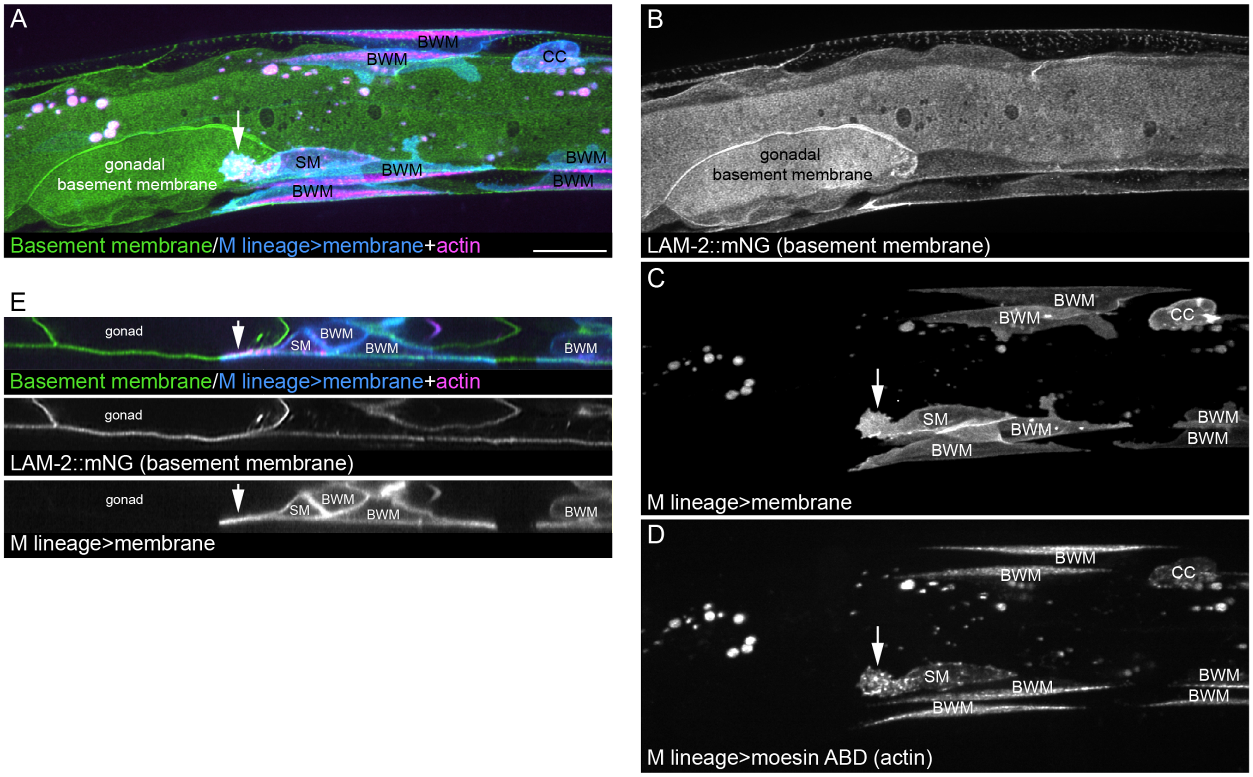

**Figure S2. ERK nKTR biosensor activity during SM migration.**

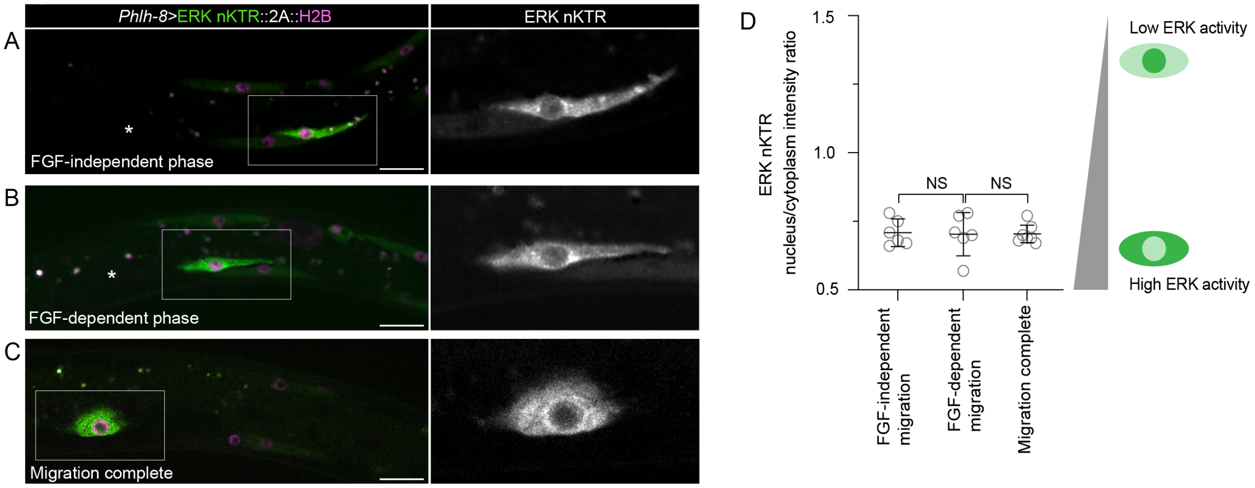

**Figure S3. Workflow for autofluorescence subtraction.**

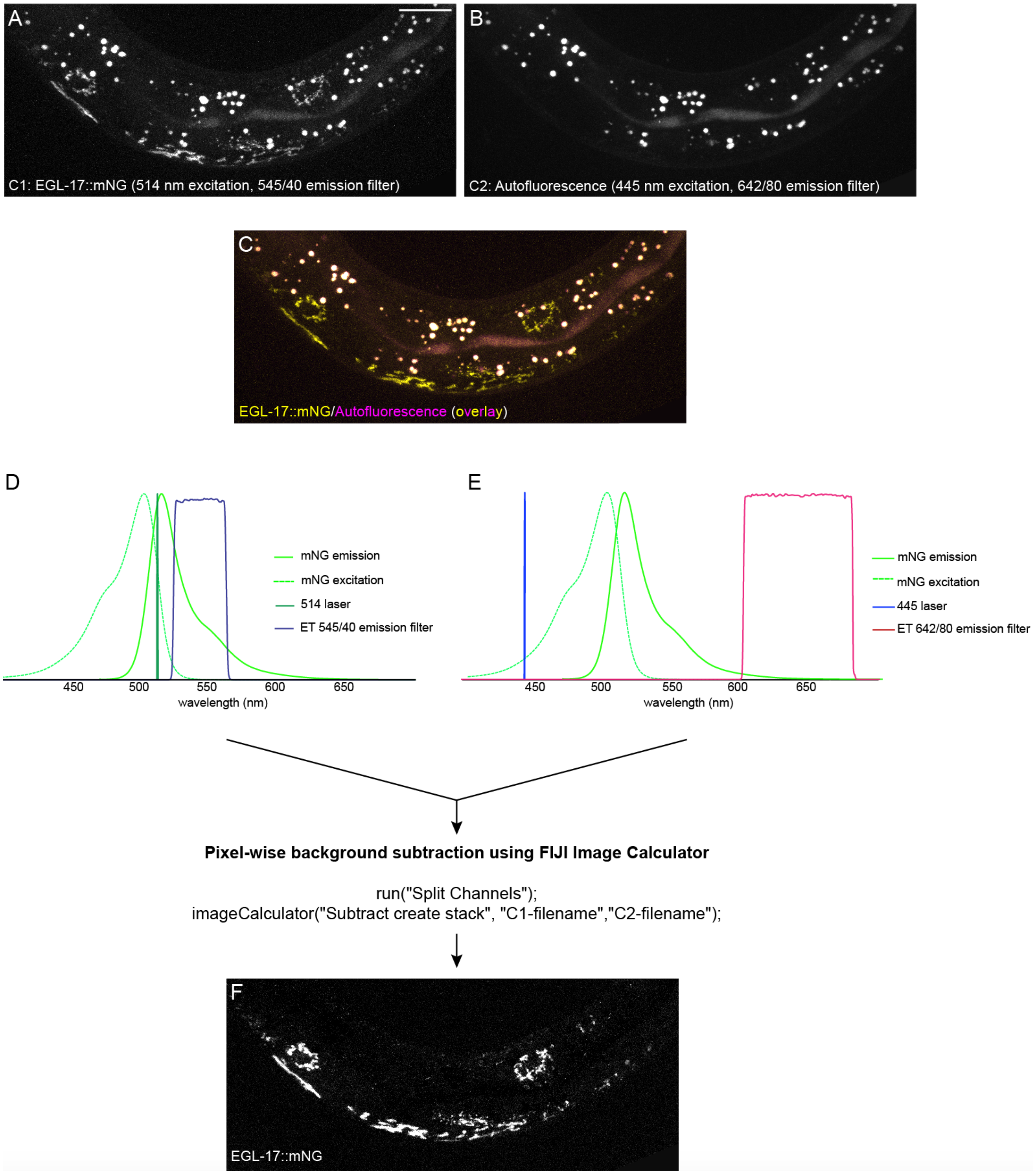

**Figure S4. Endogenous *FGF/egl-17* transcriptional reporter expression prior to SM birth.**

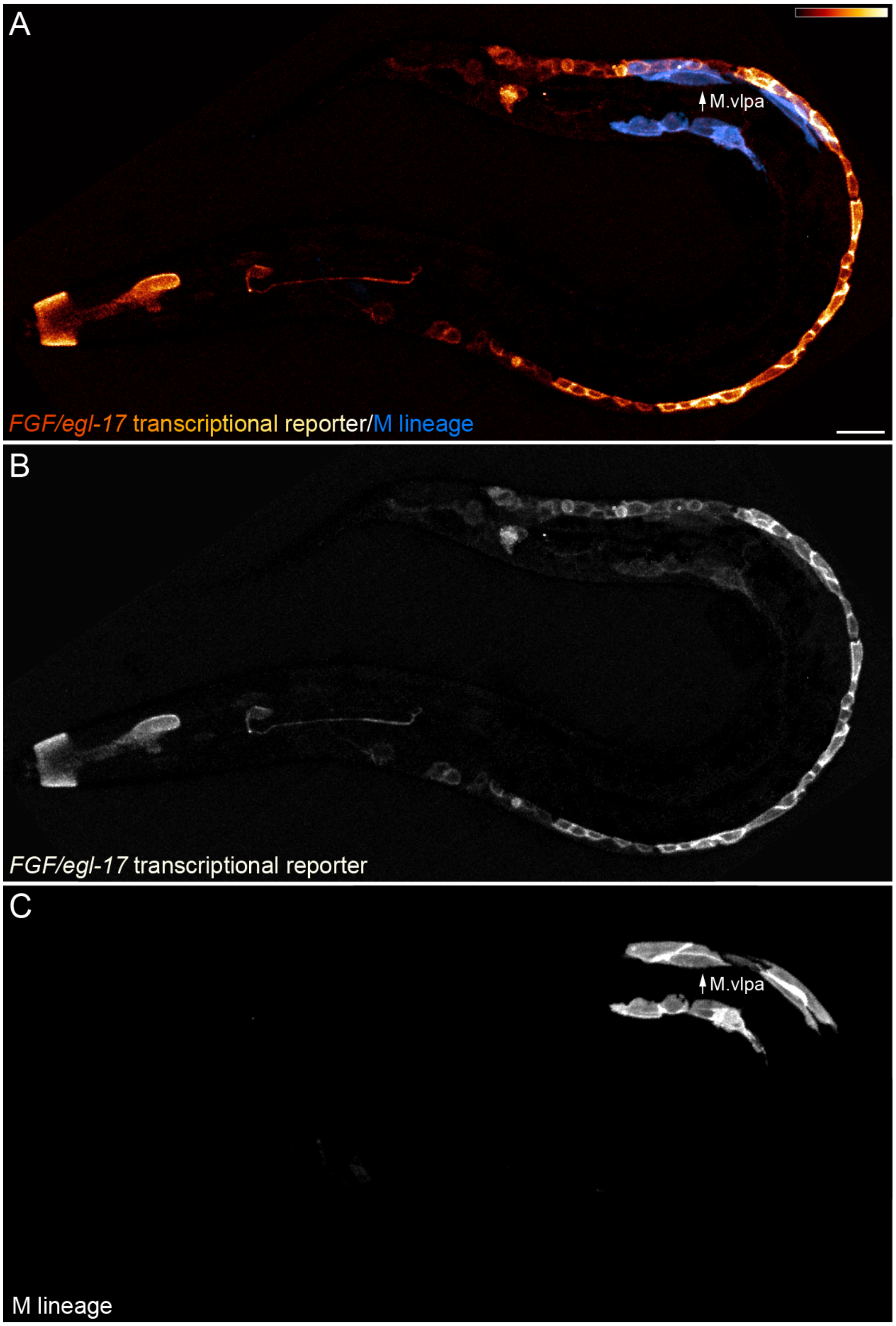

**Figure S5. Time-lapse imaging of migrating SM and *FGF/egl-17* expressing cells during FGF-dependent migration.**

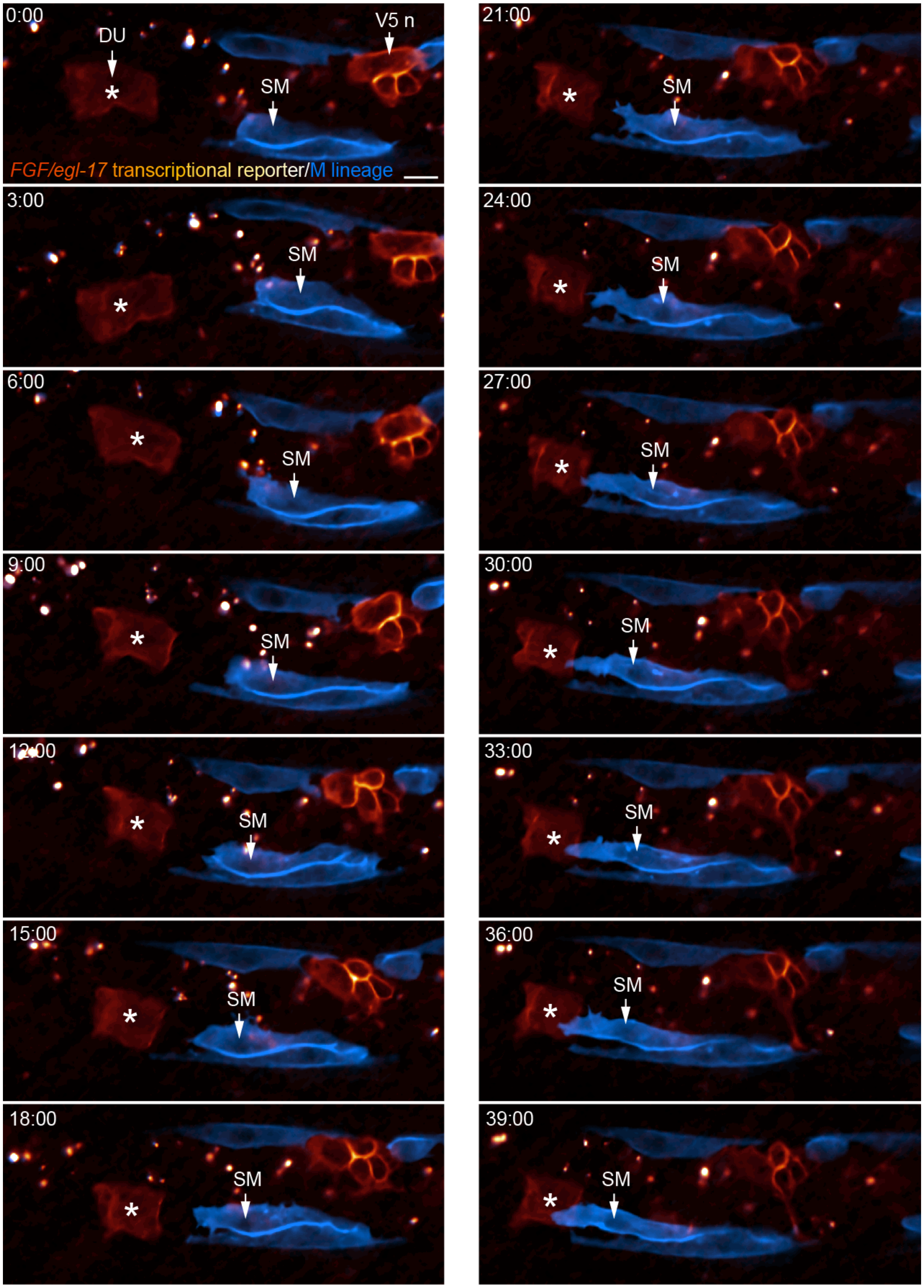

**Figure S6. Final SM positioning over the uterine and P6.p cells.**

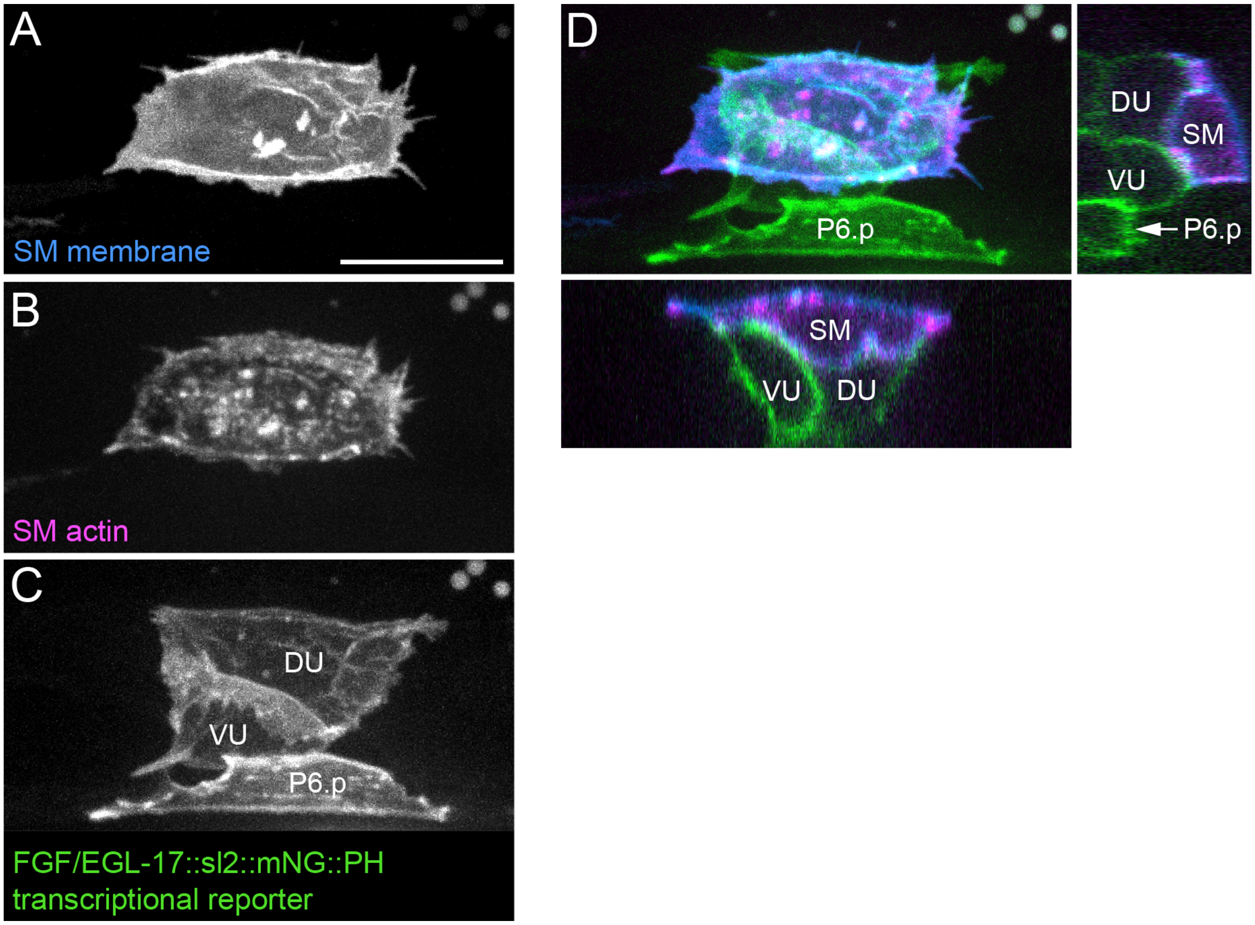

**Figure S7. FGF misexpression interferes with SM migration in a wild-type background and fails to rescue SM migration in *FGF/egl-17(Δ)* animals.**

A

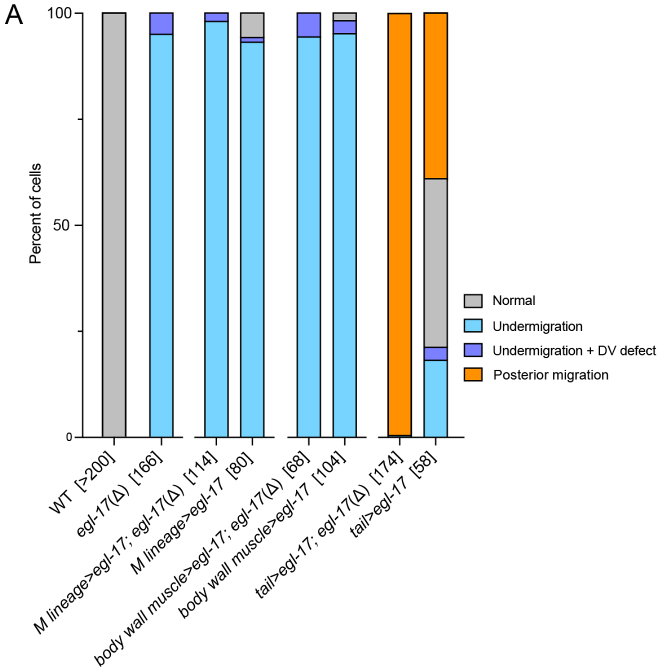


B

| Strain | *FGF/egl-17* expression [n] | normal migration | undermigration | undermigration + dorsoventral defect | posterior migration |
|---|---|---|---|---|---|
| APL5 | WT [>200] | 100 | 0 | 0 | 0 |
| APL299 | *egl-17(Δ)* [166] | 0 | 96 | 4 | 0 |
| APL201 | *M lineage>egl-17; egl-17(Δ)* [114] | 0 | 98 | 2 | 0 |
| APL200 | *M lineage>egl-17* [80] | 6 | 93 | 1 | 0 |
| APL126 | *body wall muscle>egl-17; egl-17(Δ)* [68] | 0 | 94 | 6 | 0 |
| APL342 | *body wall muscle>egl-17* [104] | 2 | 95 | 3 | 0 |
| APL242 | *tail>egl-17; egl-17(Δ)* [174] | 0 | 1 | 0 | 99 |
| APL316 | *tail>egl-17* [58] | 40 | 16 | 3 | 41 |

% of cells (scale 0–100)

**Figure S8. Precise SM positioning and normal muscle subtype differentiation in a *FGF/egl-17(Δ)* + *Pmyo-2*>FGF/EGL-17::mNG animal at the L4 stage.**

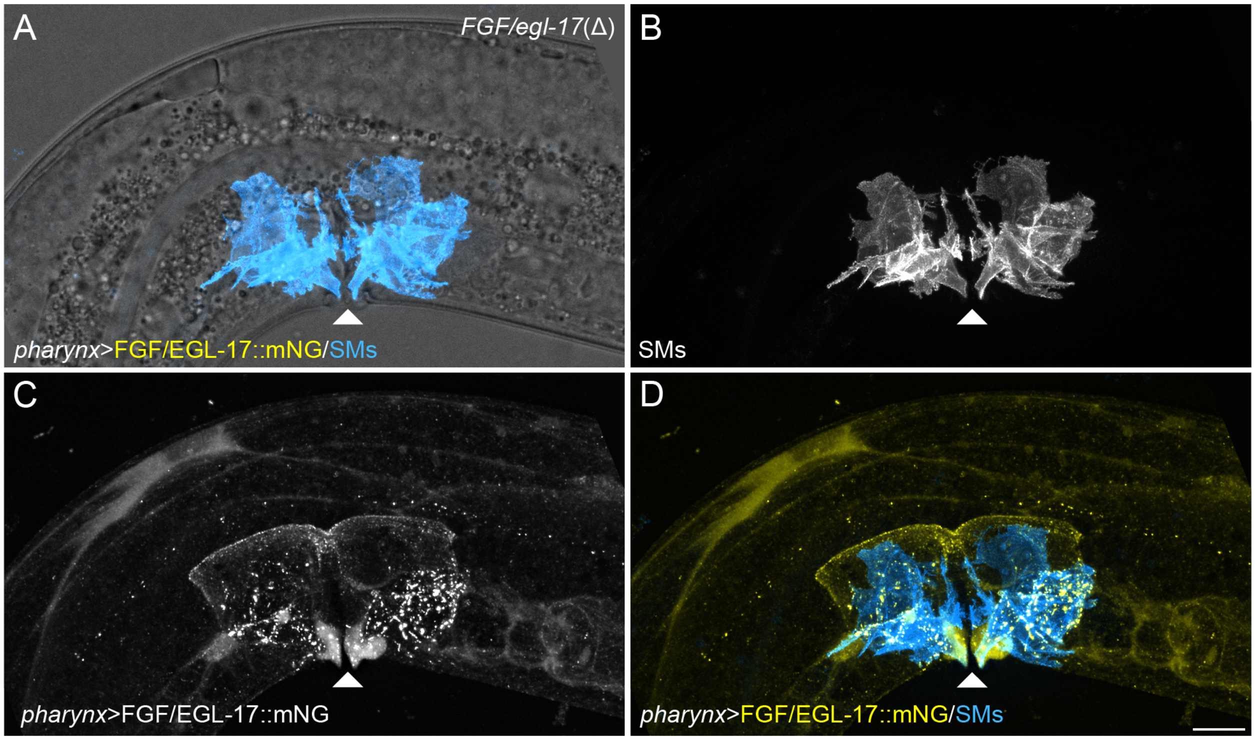

**Figure S9. Membrane-tethered FGF/EGL-17::mNG::NLG-1™ is capable of contact-dependent signaling.**

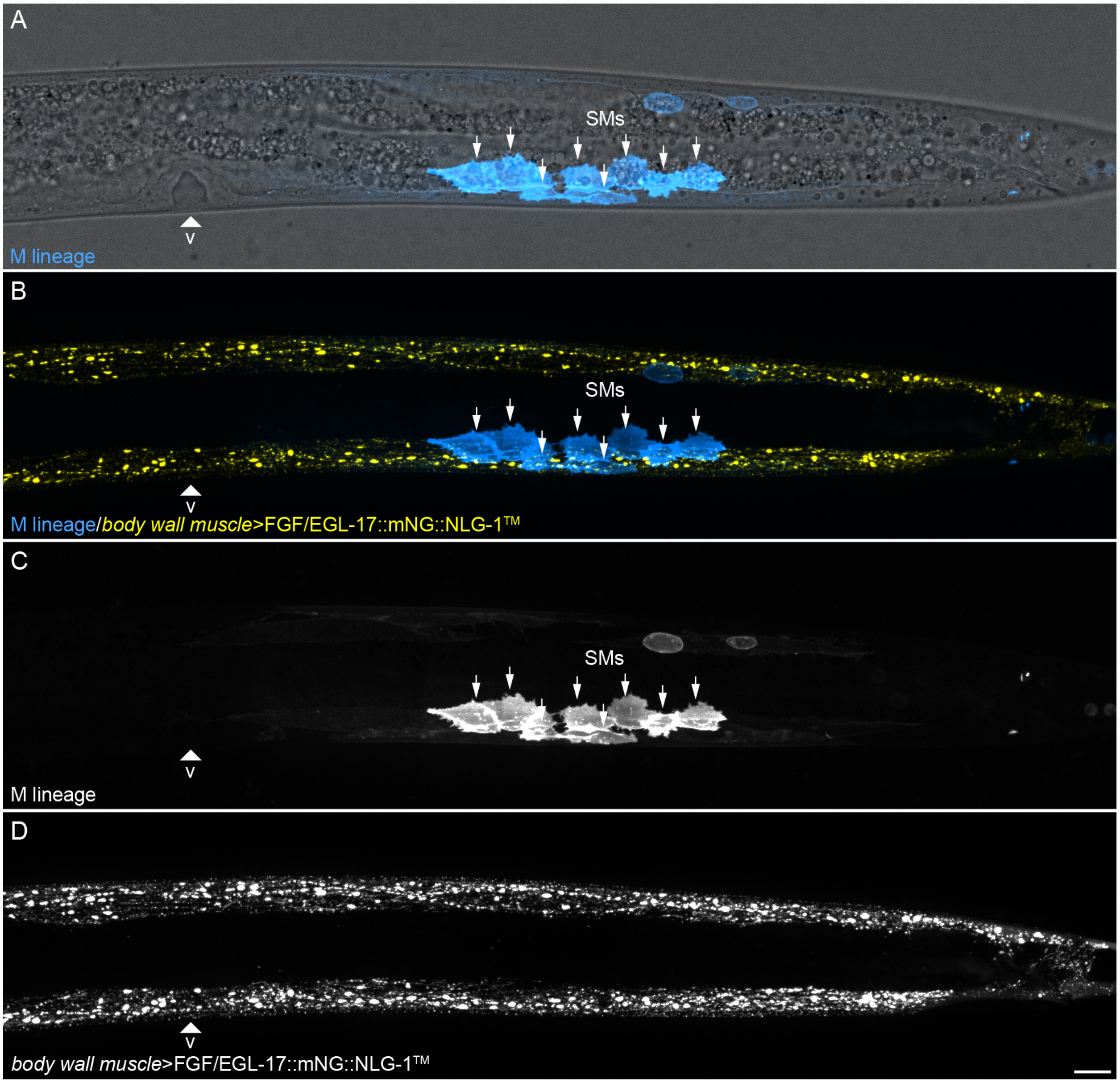

**Figure S10. Morphotrap does not sequester FGF/EGL-17::YPET.**

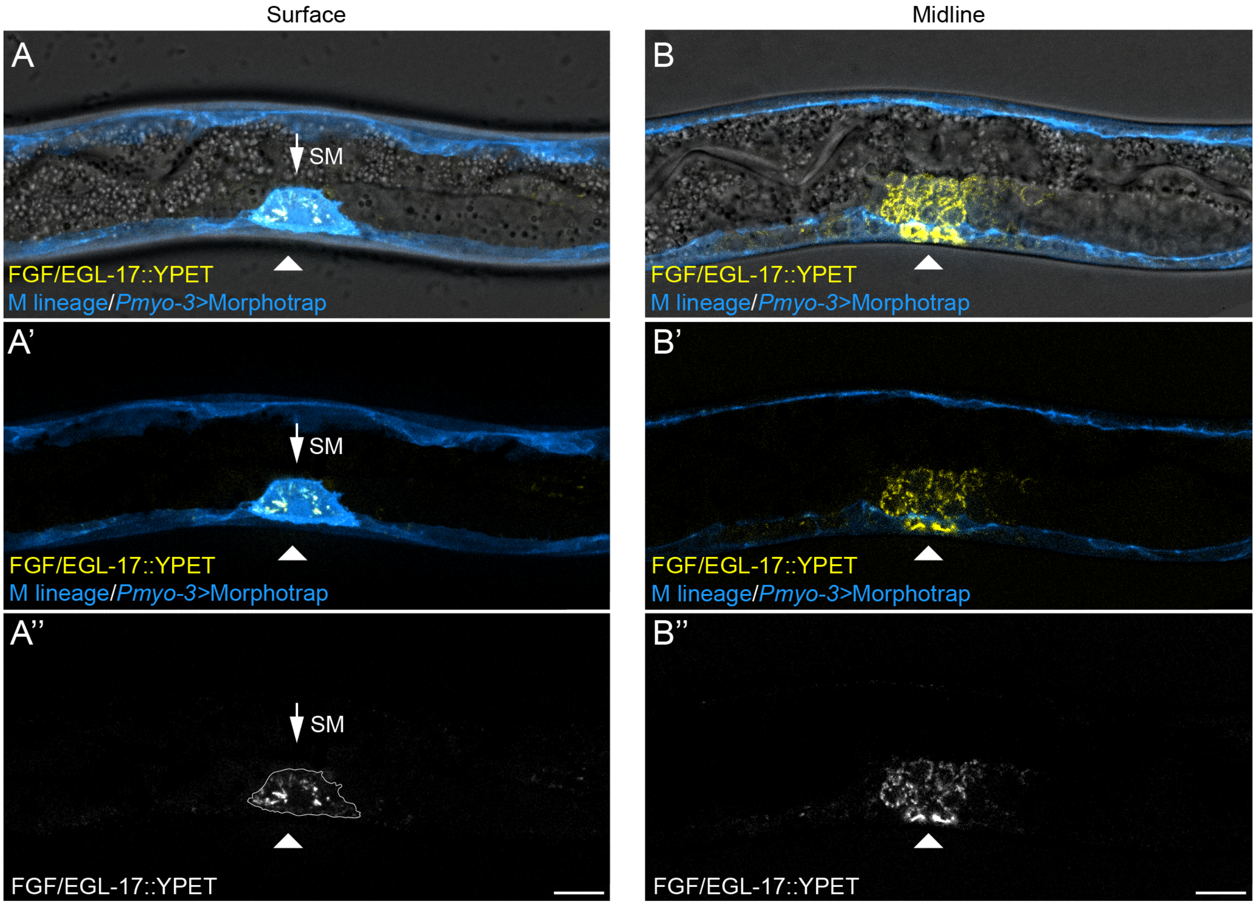

**Figure S11. Dorsoventral and anteroposterior migration defects in a *Pmyo-3>FGFR/egl-15(5a)::SL2::2x mKate2::PH* animal.**

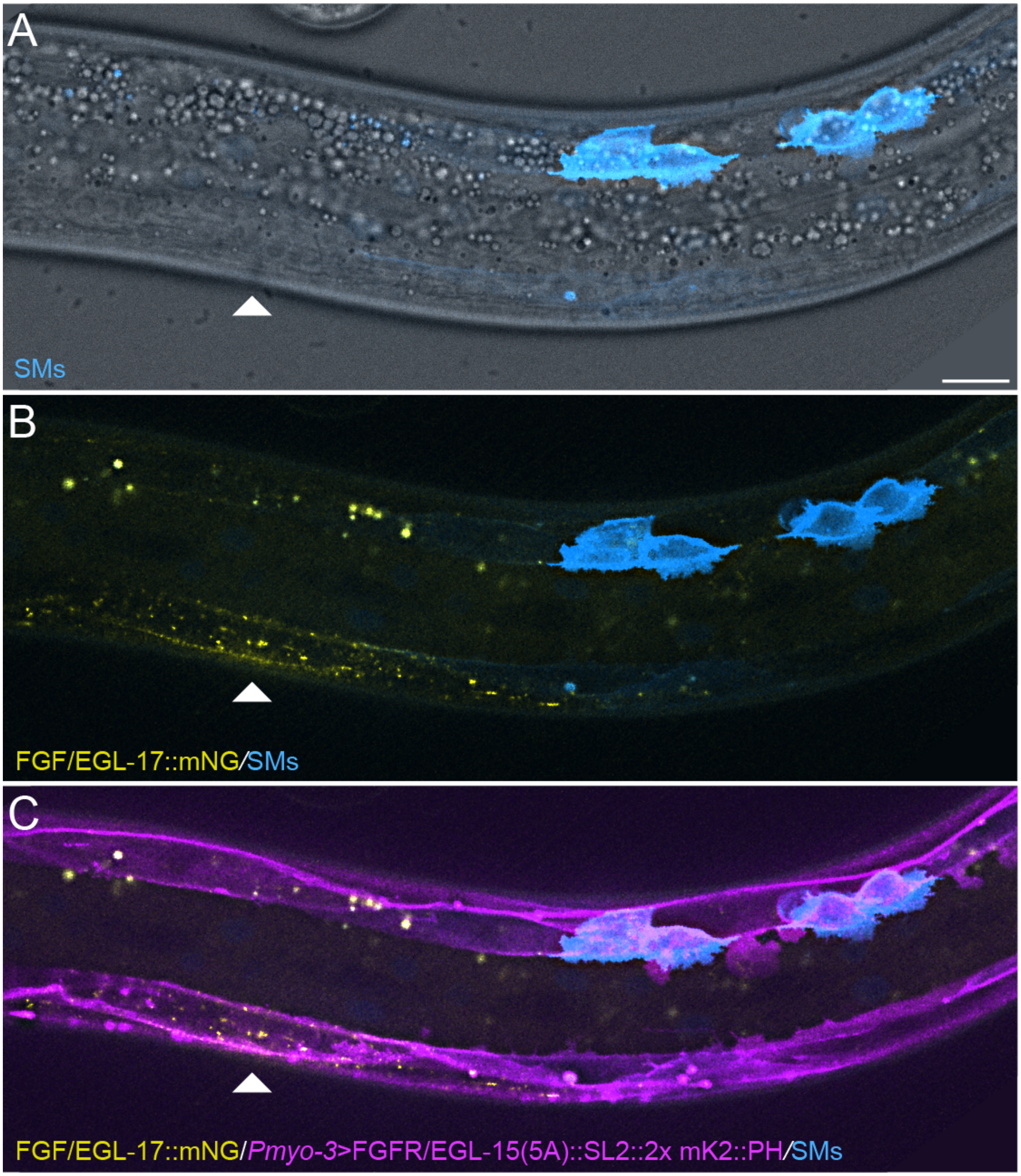

### Table S1. Source data for SM migration phenotypes

| Strain | Experimental manipulation [n] | overmigration | overmigration + dorsoventral defect | normal migration | undermigration | undermigration + dorsoventral defect | dorsoventral defect | posterior migration |
|---|---|---|---|---|---|---|---|---|
| APL5 | WT [>200] | 0 | 0 | 100 | 0 | 0 | 0 | 0 |
| APL299 | *egl-17*(Δ) [166] | 0 | 0 | 0 | 96 | 4 | 0 | 0 |
| APL201 | *M lineage>egl-17; egl-17*(Δ) [114] | 0 | 0 | 0 | 98 | 2 | 0 | 0 |
| APL200 | *M lineage>egl-17* [80] | 0 | 0 | 6 | 93 | 1 | 0 | 0 |
| APL126 | *body wall muscle>egl-17; egl-17*(Δ) [68] | 0 | 0 | 0 | 94 | 6 | 0 | 0 |
| APL342 | *body wall muscle>egl-17* [104] | 0 | 0 | 2 | 95 | 3 | 0 | 0 |
| APL242 | *tail>egl-17; egl-17*(Δ) [174] | 0 | 0 | | 1 | 0 | 0 | 99 |
| APL316 | *tail>egl-17* [58] | 0 | 0 | 40 | 16 | 3 | 0 | 41 |
| APL45 | *pharynx>egl-17; egl-17*(Δ) [92] | 41 | 1 | 39 | 19 | 0 | 0 | 0 |
| APL130 | *membrane-anchored egl-17* [162] | 1 | 0 | 0 | 95 | 4 | 0 | 0 |
| APL243 | *body wall muscle>membrane-anchored egl-17* [98] | 0 | 0 | 0 | 100 | 0 | 0 | 0 |
| APL670 | *body wall muscle>Morphotrap; egl-17::YPET* [76] | 0 | 0 | 100 | 0 | 0 | 0 | 0 |
| APL622 | *body wall muscle>FGFR/egl-15(5A)* [194] | 0 | 0 | 48 | 44 | 6 | 2 | 0 |

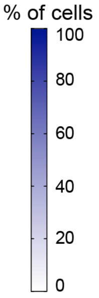

### Table S2. Resources and reagents

| Resource or Reagent | Source |
|---|---|
| ***C. elegans* strains** | |
| *C. elegans*: APL5: ljfSi2[Phlh-8>mKate2::*D. melanogaster* moesin actin-binding domain::F2A::2x mTurquoise2::PH::3xHA::tbb-2 3'UTR loxN ttTi4348] I | This paper |
| *C. elegans:* APL23: ljfSi10[Phlh-8>2x mTurquoise2::PH::tbb-2 3'UTR loxN ttTi5605] II; egl-17(ljf7[egl-17::mNG^3xFlag]) X | This paper |
| *C. elegans*: APL45: ljfSi18[Pmyo-2>egl-17::mNG::tbb-2 3'UTR loxN ttTi4348] I; ljfSi10[Phlh-8>2x mTurquoise2::PH::tbb-2 3'UTR loxN ttTi5605] II; egl-17(ljf14[deletion + mNG^3xFlag]) X | This paper |
| *C. elegans:* APL126: ljfSi33[Pmyo3>egl-17::mNG::SL2::2x mKate2::PH::3xHA::tbb-2 3'UTR loxN ttTi4348] I; ljfSi10[Phlh-8>2x mTurquoise2::PH::tbb-2 3'UTR loxN ttTi5605] II; egl-17(ljf14[deletion + mNG^3xFlag]) X | This paper |
| *C. elegans:* APL130: ljfSi10[Phlh-8>2x mTurquoise2::PH::tbb-2 3'UTR loxN ttTi5605] II; egl-17(ljf25[egl-17::mNG::nlg-1 C-terminus]) X | This paper |
| *C. elegans:* APL199: ljfSi10[Phlh-8>2x mTurquoise::PH::3xHA::tbb-2 3'UTR loxN ttTi5605] II; egl-17(ljf24[egl-17::SL2::mNG::PH]) X | This paper |
| *C. elegans:* APL200: ljfSi34[Phlh-8>egl-17::tbb-2 3'UTR loxN ttTi4348]) I; ljfSi10[Phlh-8>2x mTurquoise2::PH::tbb-2 3'UTR loxN ttTi5605] II | This paper |
| *C. elegans*: APL201: ljfSi35[Phlh-8>egl-17::tbb-2 3'UTR loxN ttTi4348]) I; ljfSi10[Phlh-8>2x mTurquoise2::PH::tbb-2 3'UTR loxN ttTi5605] II; egl-17(ljf14[deletion + mNG^3xFlag]) X | This paper |
| *C. elegans*: APL242: ljfSi32[Pegl-20(-1261 – 610)::pes-10>egl-17::mNG::SL2::2x mKate2::PH::3xHA::tbb-2 3'UTR loxN ttTi4348]) I; ljfSi10[Phlh-8>2x mTurquoise2::PH::tbb-2 3'UTR loxN ttTi5605] II; egl-17(ljf14[deletion + mNG^3xFlag]) X | This paper |
| *C. elegans*: APL243: ljfSi36 [Pmyo3>egl-17::mNG::nlg-1 C-terminus::tbb-2 3'UTR loxN ttTi4348] I; ljfSi10 [Phlh-8>2x mTurquoise2::PH::tbb-2 3'UTR loxN ttTi5605] II | This paper |
| *C. elegans*: APL299: ljfSi2 [Phlh-8>mKate2::*D. melanogaster* moesin actin-binding domain::F2A::2x mTurquoise2::PH::3xHA::tbb-2 3'UTR loxN ttTi4348] I; egl-17(ljf14[deletion + mNG^3xFlag]) X | This paper |
| *C. elegans*: APL311: ljfSi41 [Phlh-8>2x mTurquoise2::PH::3x HA::tbb-2 3'UTR loxN ttTi4348] I; ljfSi42 [egl-17>2x mKate2::PH::3xHA::tbb-2 3'UTR loxN ttTi5605] II; arTi133 [Phlh-8>ERK-nKTR::mClover::T2A::mCherry::H2B::unc-54 3'UTR] | This paper; De la Cova, et al., 2017 |
| *C. elegans*: APL319: ljfSi37 [Pegl-20(-1261 – 610)::pes-10>egl-17::mNG::SL2::2x mKate2::PH::3xHA::tbb-2 3'UTR loxN ttTi4348] I; ljfSi10 [Phlh-8>2x mTurquoise2::PH::tbb-2 3'UTR loxN ttTi5605] II | This paper |
| *C. elegans*: APL342: ljfSi38 [Pmyo3>egl-17::mNG::SL2::2x mKate2::PH::3xHA::tbb-2 3'UTR loxN ttTi4348] I; ljfSi10 [Phlh-8>2x mTurquoise2::PH::tbb-2 3'UTR loxN ttTi5605] II | This paper |
| *C. elegans*: APL622: ljfSi2 [Phlh-8>mKate2::D. melanogaster moesin actin-binding domain::F2A::2x mTurquoise2::PH::3xHA::tbb-2 3'UTR loxN ttTi4348] I; ljfSi39 [Pmyo-3>egl-15(5a)::SL2::2x mKate2::PH::3xHA::tbb-2 3' UTR lox511i] IV; egl-17(ljf7[egl-17::mNG^3xFlag]) X | This paper |
| *C. elegans*: APL670: ljfSi40 [Pmyo-3>pat-3 signal peptide::2x vhhGFP4::CD8 transmembrane domain::2x mTurquoise2::PH::tbb-2 3'UTR loxN ttTi4348] I; ljfSi10 [Phlh-8>2x mTurquoise2::PH::tbb-2 3'UTR loxN ttTi5605] II; egl-17(ljf27[egl-17::YPET^3xFlag]) X | This paper, Pani & Goldstein, 2018 |
| *C. elegans*: N2: wild-type | CGC |
| **Plasmids** | |
| Plasmid: pAP082: Peft-3>Cas9 + PU6>Chr I (ttTi4348) sgRNA | Pani & Goldstein, 2018 |
| Plasmid: pAP087: 2x mKate2::PH::3xHA::tbb-2 3'UTR loxN SEC loxN backbone for cloning transgenes, Chr II | Pani & Goldstein, 2018 |
| Plasmid: pAP088: 2x mTurquoise2::PH::3xHA::tbb-2 3'UTR loxN SEC loxN backbone for cloning transgenes, Chr I | Pani & Goldstein, 2018 |
| Plasmid: pAP092 Pegl-17>2x mKate2::PH::3xHA::tbb-2 3'UTR loxN SEC loxN, Chr II | This paper |

| | |
|---|---|
| Plasmid: pAP137 Phlh-8>2x mTurquoise2::PH::3xHA::tbb-2 3'UTR loxN SEC loxN, Chr II | This paper |
| Plasmid: pAP138 Phlh-8>2x mTurquoise2::PH::3xHA::tbb-2 3'UTR loxN SEC loxN, Chr I | This paper |
| Plasmid: pDD122: Peft-3>Cas9 + PU6>Chr II (ttTi5605) sgRNA, Addgene #47550 | Dickinson, et al., 2013 |
| Plasmid: pDD162: Peft-3>Cas9 + PU6>empty sgRNA for cloning, Addgene #47549 | Dickinson, et al., 2013 |
| Plasmid: pDD268: mNG^SEC^3xFlag backbone for cloning repair templates, Addgene #132523 | Dickinson, et al., 2015 |
| Plasmid: pDD283: YPET^SEC^3xFlag backbone for cloning repair templates, Addgene #66824 | Dickinson, et al., 2015 |
| Plasmid: pDD315: mTurquoise2^SEC^2xHA backbone for cloning repair templates, Addgene #73343 | Dickinson, et al., 2015 |
| Plasmid: pRCA5: Phlh-8>mKate2::Dme moesin ABD::F2A::2x mTurquoise2::PH::3xHA::tbb-2 3'UTR loxN SEC loxN, Chr I | This paper |
| Plasmid: pTG42: egl-17::mNG^SEC^3xFlag::nlg-1 C-terminus homologous repair template | This paper |
| Plasmid: pTG003: egl-17::mNG^SEC^3xFlag homologous repair template | This paper |
| Plasmid: pTG004: Peft-3>Cas9 + PU6>egl-17 C-terminus sgRNA1 | This paper |
| Plasmid: pTG006: egl-17::YPET^SEC^3xFlag homologous repair template | This paper |
| Plasmid: pTG009: Peft-3>Cas9 + PU6>egl-17 C-terminus sgRNA2 | This paper |
| Plasmid: pTG012: Peft-3>Cas9 + PU6>egl-17 N-terminus sgRNA for deletion | This paper |
| Plasmid: pTG034: egl-17 deletion + mNG^SEC^3xFlag homologous repair template | This paper |
| Plasmid: pTG052: egl-17::SL2::mNG::PH^SEC^ homologous repair template | This paper |
| Plasmid: pTG077: Pmyo-2>egl-17::mNG::tbb-2 3'UTR loxN SEC loxN , Chr I | This paper |
| Plasmid: pTG110: Peft-3>Cas9 + PU6>Chr IV sgRNA | This paper |
| Plasmid: pTG125: Pmyo3>egl-17::mNG::SL2::2x mKate2::PH::3xHA::tbb-2 3'UTR loxN SEC loxN, Chr I | This paper |
| Plasmid: pTG238: Phlh-8>egl-17::mNG::tbb-2 3'UTR lox511i SEC lox511i , Chr IV | This paper |
| Plasmid: pTG282: Pegl-20(-1261 – 610)::pes-10>egl-17::mNG::SL2::2x mKate2::PH::tbb-2 3'UTR lox 511i SEC lox 511i, Chr IV | This paper |
| Plasmid: pTG284: Pmyo-3>egl-15(5A)::SL2::2x mKate2::PH::tbb-2 3' UTR lox511i SEC lox 511i, Chr IV | This paper |
| Plasmid: pTG327: Pmyo3>egl-17::mNG::nlg-1 C-terminus::tbb-2 3'UTR lox511i SEC lox511i , Chr IV | This paper |
| **Key Reagents** | |
| Gibco Hygromycin B (50 mg/mL) (Fisher Scientific) | 10-687-010 |
| NEBuilder HiFi DNA Assembly Cloning Kit (New England Biolabs) | E5520S |
| Invitrogen PureLink HQ Mini Plasmid DNA Purification Kit | K210001 |
| Q5 High-Fidelity 2X Master Mix (New England Biolabs) | M0492L |
| Q5 Site-Directed Mutagenesis Kit (New England Biolabs) | E0554S |

**Supplemental File 1. Molecular cloning information.**
This file includes sequence information for transgene promoters used in this study, the *nlg-1* C-terminal region used for membrane tethering, and a Chr IV transgene insertion site that has not been described elsewhere.

***egl-20***$^{(-1261 - 610)}$ **enhancer:**
<u>*Underlined sequence</u> is the *pes-10* minimal promoter.
gttggatgttttcttataaaattatcttggaagtttttttaaaaattaaatactctgttacatttttttgcaatgttt
ggaattgaaatatagaacaatataaaacttctagagagaaaaattacttccagaatattgataaatgataaagaata
ttgataacctgaggatttgccactttttgacggcagcattaaaaattgcaattggatactcaagaagataaagtgat
caattatggagaaaataaaatactgtgtaaataaatgatttttttaaacatttttttccgaaatataggtgttccat
ttgtcacatgataattatgaatattcttgaatttctatgttttgaaaaattgaagctaaaaaataagtgtcaaatat
tgttaaagtcccaaaaatcttggaatttttcgtttcgtattgaccattcacttcttcccatcctttttcaccatttc
aacgtcttcctcttctatttgcataccgacaacgtcttcttctcttccattttctgaatagatccacgggccataaa
gcatgaattggaaaggatggagaagatgacaagataccacatccagcaccatcaattcgaccttcgaattttctaaa
cacat<u>tgacattcctcttattcatcattcttcatccctgcaggatcgatttttttgcaaattacgagcgttgtagggg
gcggagcgataggtcctataggttttggtatatcatcattcattcattcattggtacattcatttacccaccttcct
ctttctgagcttctctggagttctgtgcttcctttttcccttatctttatactgtaatttttaactttcaggcattg
attggatcattggccaaaggacccaaaggtatgtttcgaatgatactaacataacatagaacattttcaggaggacc
cttgcttggaggagctcagaaaa</u>

***hlh-8* promoter:**
aaagctttcaaaatcccaaacatagcatcttacttactcgcctttgctcccaaagatttaccattttcttgtttttg
tttgttaatttcctctcccacccattttcaaccatgtcatttcacacattaaactcagtatgcatcgatttaattgt
tccaggctaattgggtttctgtcattgttaatagacacaatttttactgagaaatccttttcatatgtgtgaaattt
gttctttaaacgtgcatttcctatttatcgctctctagatttgatgccagttcgtgtagtttttttcccaaccgttga
ctcatcgataccgcaatccatttcacccagtatttgtaccattgccgtgaattcacagtattacatggtcgtcccct
tttccacaaaatctctttcagactccctccaccctgatcccgttttaattgtttcttgctgatttccctttcaaccg
atcttcgtttttttttcaatagggtatatttgaaatgtcggaatgtcgattcagcagtaaatttgatatgacactct
gactttttttaaatgtaaatctgattgtttatcagaattcaaccgggcccactgattattcattaattatgcgacaag
aagaataacgttatagtttttctttcttaagcatttttcatcatcaacatttttttgtttagatttaattttgttgct
aataaaaaactacacaaaacgccaagcacatttatttgactttttttttttgagtaatgcagtaagtaggcaatttgt
tgtttttataattgattagttttaaggcctacttgcaaaactgagaaaattcttatcaagaattcaaaaccatttac
gattatcggattttttaatatggacaatgttactcattttggataaatcaaaagagtttttggattttaaaattaata
actgtttattactgattaacaaaagattacggtaaattggagttacaaaatggacaagttttatttatgaatcaaga
catattttgataaaaaatatgggtacccccgccaatttaaaaataaatttttttaacttaaaaaaaaacgtgtgaaat
gctttcatattttcaggcttacttagaaaaacaccattcctaagtctaacgaggaaaatgggaaacatgggaaatat
taccgaaacctgggaaatatttttttattgattccaaattttcccttgattccaaatatcgatgtgaaaaaaaattaa
aacaaaaattactgattttatttaagcttgaaatcacaaattccattttttatgtcatacttcagatttaaacaaatt
tattttatgtgtgtttttttaatggtatgtcctaacgattttcctaattgacaactattatagattgaaaacacaga
atgccaagtaacgtaagaaattttttttttcgaataccagacataatttcaataaaagaaaaatctttaaaaaaagtt
taactttatactacaataattttggtttttatgaaggaatctgtattgcggcacatcatgtatttgctcaaggtttc
gttggttgacatattttttgtgatccgtggaatgagcttgcatagagtttagtcataattgggtttattttacataca
gtttgaagtatgttttgtctgttattcgaaattttttagttagtttggtaagttgaaaagaatttacaaaatttaac
taaacgaggacgctcatctcgattcttataattccataataattctacggtaaaagtcaagttatgcctcaaaacat
gtaatacataattaccaaaactacttaattaccgcatttccttagtattttaatagtggtgtagtcgaattttttttt
attgctttattagactcaaaattgtctgcaaacacccaatttcataatgaaacttattgaaaacaatacactttgaa
acaatttagtatttcaggaataaggtcactggaacttcgaaaatgataatttgaataaccgttaataaaatattcaa
accaatttgcataatcttgattttgtatcatgatggtatagaatgggacttttgaaagatgtgaagtttcaattcag
taagtacaactttaaaattgggctgcaagatttcttcttaaattttcaacagtttcaaaacactgaaatcatgctga
actatggttagagactcgacaactttcctaaattttggagcagtttcaatggttttcgaatgtatactacatttgta
ccattttatgtgttagtgtaggttgctttgcttctaccttcactctcaaattctttttttccgcggtaatttttcaac
tatcgaaacaaagaagaagtcatgaataactgtccgcatagagttctcacccgtcccctttcttttcaccgagcagt
cacgagagaaagaagaagaagcggaccgctgcagagattcttcgtagcggagccccgcccatcctctgttacgcata
tgtttggatctcatttcttccactcattaacgttccaggtatgtttttttttgttgatttttttaatgtgttacttttc
atatgtttattacttgaactgaccgatttcaaatatgattttcacag

***myo-3* promoter:**

cggctataataagttcttgaataaaataattttcccgacaaaacatgagtatttctttcgaaaataaaagtgcaggc
taattagagattattctgtaattaactgcataatttgtcacgtgccatagttttacattccactacgtcatagttct
taaaatactaatctcctgaaaatagaagtaggtgaagaaagtttaattatcagttctaaaatgacaattgatctttg
gaatatgttctgaaactaccgatcattgaacagatgctatttgaatgatatagaattgtatatttgcaatttctgaa
acgcgttcttaaaggcacacagattaattcaaaagggtctggccgcaaaaaggtttatggtggccgattttgagttt
tgtgtgtgattgctttttcacaatcagtgttttcaggattatgtgatgaactagatcttcaagtttcgttacatttc
atatgttttcggaactcacgaagtacatattgggtattgtgctcaaaaaattcagcaatcagcttcgctccgctgac
tttagaacccaaaaaaatagtatggccaaactgactgtgttacgatcatttcaatttttcaatacatatttaagatt
tctaagagtaagaaggtcaaaaactgttctggaatacatatatatttttcaggttacaattagtcaaaaagtgcact
gaaatatacgttttaatttcacgaataacccaattagttcaatgtatttttggtcaaccaacgttaaagtttggctt
ccaaccaattatcatttctgatcaaccacaatgtttttctttatctgcaagttaatttttattttatccagatg
tttggcatatttttcaattcttcactagcgcccacttcttgcacttccggcgccctgaatctaatgcatctgttgca
agaattgaaagaccaatcaacacattgttttcttcacgagatactgaagaaaatgaataaaaacagagaaaaagagc
catgtgattagtgacaactgttgctaacagataccatagcttggacttggtacgtgatggcaacgtatgggtcaaca
aaaatgattgcagagggggtgcaaaacagtcaagtcgagaaaatatgaaaaacagaaaacaaagaacagaaaaatgg
gtttgagagtcagtataatttataaaagaaaaattgtacatagaaattaaccattttgtagaagaagttatttttc
aagcatcgttaaaaattattcaaagcaccttatttcatatttaattttaaacatggttaaatgaacaacacggtgcg
caatcaggaaaacttgaaatctgaaactgttgttgtgatcttcttcgcaactgttcagatagcactagtgtaatgtt
aagagtgcgcgaatataatggaatataatggatcacacctcctgccatcaggtaaacgtctctgttatcacatattt
ccaactattaaattttaccttttacagttttacatttttttgaaaaaagtaacttttttgtcttcaaaatccctgac
gaaaatatcaaatattttaatcgagactgcagaggaaccgattgatgatttggaaaatccagctttacctgtgtaag
aactgaaaagtttcataaccctagggtattcccagttacattccccactggctaacaatagcacccagtttttcatc
acctttcttcaaatttctcggcgatttgttaaaaacaaaatttgtgtcccttctctgatatctctatgtctctaaac
acaagttcatcggaaaacgaaggagggtaggtgttggttgggctcccgaagtgaaaatagaagagcaagaatagaat
attagagagagagtgcagagagggcgggatagctcccgggattccgttttcttcttctttatcttcaacgatgatgt
gtgtgcgtgttgtatagattctgttgctcccccacaactcgctccgaaggctcaatacaattcaattgatattggag
gagagcctaccggagtgggaggataagaagaaacataagaagaagaagaagaagaagcatgcttctggtttttgatg
ctatgaaaacggcacaaaaagatgattgaggtccctttcaataccttctctcatctttcaaatcccattgaaacct
aaaacttctcaccacgctttaccattgttctccaaaaacttatagcaatgtctataacttttttatctctgaaaagc
agtgttccatttttctttttcctattttatttcaattgtttctcacatttcgtttggattctttgcttgtcaaccag
cttcttcttccacttttaccgtctaattttcagggcagggagccatcaaacccacgaccactagatccatctagaa

***nlg-1* C-terminus**

Transmembrane domain indicated in bold. Exons are capitalized, introns are in lowercase.

TTGCTGTCAGATCATTTCCGAAAAGACTCTTACTTTGGAAAAACCCGTCATTTTTCTTCATATGCGAATCTTCCGTT
CCCGCCACCACCAATGCCCCCATCACCACCACCAGAGCTTACAACAAAGCCCAAACCAAGTGAATgtaagttttgat
agttaagttcaaacaaatctagttgtataaacaatacaagttttagagttttagacaagtgagatttttgagaaaaa
ataacactgctataaaattcttaattaaatgccttgactcgtttttgatagtcgaaaaaaagtcacaaaatactaca
aaacctctagaacccagttttattaagacggatattgaaagcttgttctacaagtttttaacagtgctttcttgtt
ttttccaaatattttctaactctgactggaatttgaaaaaactagttgatattttgaaataacgtttcacgttgaaa
aatatttctagtttcaaacatttttgcccttttgaaataaataactatttcatattcaaacctgccagcttccaccg
gtattaatactccacccgataagttgttttctcatccgaaactagaagaacctaagctgtttgaaaacattattgta
attagaaatgattcagCGCCCACAACTCTCCAAACAACAACTGAAAGTGAAAAGGCCGCCGCTGGAAGTTTTACTGG
AAAAGC**TCTCGGTGGCGTTATTTTCATCGGTTGTGGATTCCTCATTATGAACGTTTGCCTATTAATTGCTGTTCGTA**
**GAGAA**gtaagttttctgctttgccttcccttcagtagtagttccagtattgttgtcttgaactgtacacgctggtga
tcatgacaatggagtagctatctatcttattcagttgaatttttgtcacttgattaagttattttcctgttacacat
tatcgatatttag**TGGGGAAAGAAGCGGCGGAA**CGAGAAGAAGTTTCAACTGCAGTATCAGACTTATAACTCCAACC
ATGGCGGCGGCGCGGAACAATACAACAGCTTAAACTCGCCGgtatgaactatttgcttctcttttgtgaatttgttt
ctagtttttttttctcgattgatttgaaccaaaaactaacgaaattcaactagcttaacctttcatccaacgcacgag
catggctcagaactcccccctaatccttaatttggttatcagttatccctcatgccaccacctccgccaccgctcaat
ggtgttcacgacgacattttcgatcatcgggtgccgcatcttcggaacggatccacggttcacggcacgcttccgag
gcattcatttcaggaacaggcacgtgtataatgcattatcctctcaccctgtttgaaaatctttttttcaagttatat
ttttctctcgctttttggtatagatttttttccaatttttgaagttgaagtttccaaatggtcgtagaacgaatttgt
tagaaaattgtgaattaataataattaactttcgaaatatggcaattttaacagctcaaaagtaagaaaataatgta

gtttaaacttaattatgatataacttgccaaatgtaattttgcattttctgactttaaactgataacttattagatt
tttaagcattcattgggcccaatcaaagatttgaaaaattttacagtgtttaaataaatttctcgaaatgagttgac
tcaatttagtagaaaacataatcgattgaaatacaaaacaagcgaatctattctaaaagcactttcctttttttaaa
tacctttcttttaaagttcatatttcaatattcaaataattgcccagttttaaacgagtgtgacattttgaccgttt
cgcacaagttagcagtcacctttcaatttttgcaattcactaacaaaacttgttttgctaccatttgatctaacttc
gaactaaacggaatgacctacctcaacctgcgctttctgatattcataaccttttcattaaaaaataaatgttgcgt
cccttgtccattcagttcttgtaattgtgagatcagtcgaaaacttgtgtgcttttagGAACCCTTACTATCCGCAT
CGCACAAGAACTCAACTTCGATGCGACCCGCGGAATATCACCAACGTGTCCACGTCACGGACGTGCCGCGCTTGCG
CTTCAAAATAGCCGgtaagtgattgcagcgtgttcttttcatactgacatgttcaagAGGTAACAGTTTGACTGCTG
CTCAAGCACCGACATTGGAAGAGATACAGGTC

**Chr IV transgene insertion site (Chr IV:4,237,723)**

**Guide RNA:** ACTGTTGGATGCCTGTGTAG

**5' homology arm (Chr IV:4,236,614 – 4,237,722):**
aggaatcttcttcgcggacggaactcgccgctaagcggatcgcctttaatttattctttgccttatgcgttgctcaa
aataaaacattatttcatccaaaataatttttttgaggttttagatgaatttgatagtttagcttgtcactttctcct
gtgcgaagaaacttgctaagcaggtaagcagcatccttatcaagtttcgaccttatctctcagtagtcgcatacgtc
gtatccctcatctgctttgttttagccacttcaagggaattcagatcactttttctcatgtgttttcaaatttactt
ttgtggagtgaccttaattgaccctctcgtcataagtggacggtgggtagttacttaacaaatttcgagtacaggat
agtgtcaatggcatgagacatgtagtagccaagcaattatggcgaataagtttcaaaaaccaacattctgtcaaata
caaagacctgcaacattagagaattcattaaatacatcttcgaggaaatcaaattttcccgtagacgatcccgtaga
cctacttgtacaatatgttctagatatacactccgaacttaaaatatctcgaacatttgtgttgaatatttatttta
attctaggtttttcaaattccgcacgatttttgccatactactgcccaatagtttacatttattgggcctggatctg
taaggcacttttgggagctccctttcttttccttcaaattacccatacagatctataaatcacaatttaatacaaga
ttttattcaagttaaaaacataaaagggggctttgcggtggtccccatttcaccagagaactataaaaaatataaatt
taacatcaaatgttgcaattgattccctctcaaagctgcttgaagaaccctgattctgtcaagcctatgaagattta
aaaaaaattgggaagacccttagttccaaacaagtgtcgttgacccagtaggcgatattctgaaaagtcataaaatg
gggttgctaaaaaattggtcgctaacttacatttagctaggaatgttaattggaataactcataatttacagtaaaa
ttatcaaaaaatagcaaaaatcaatagagga

**3' homology arm (Chr IV 4,237,729 – 4,238,654:**
cacaggcatccaacagtacgagttctgcgtttttgaattatagaatcaagcatgctccgcgtttgctgtactatttc
tcgcagagcagtttttttttctcacactttattattctgtactgcccctccacgctataatttttttgcatagagggcc
ataaaggaaaacggtacggtaaaaataatgggtcctttttaccgttgtgtagattgtttctaacgggatgcaaattt
tcactaaaactctggaaaactgatggaattctacgggcagccggcaattttagatatttgaagaatccttcttgatc
agtgccaagctttcccatggaccaaagatgtggatccaagatcaaatgcacaggatctgaactcctttgttccaaaa
acaaggatttcttttctatagtttgacctaaatattatagtagaacatatacatttgatgatttcatgaataattta
ttgttaaaaaataatacatgtgttaaaaaataattctaaaaaattaattggagcgtccttggcggatatggcggatt
ggaaggtcataagtaactcgaactaacctgtaaatataattaatttagaagtaatcattttgaaatgtttctaaatt
tttgtaaacgtaatcaatgatttacttacaaaatttttcgaaatattagtagaaatataaaaaaccagtaaaattca
aaagaggccagccacacatgcatcttacaatacgaatgatgcgtttttttgtcgcctcggagaaggaatcaagcaggc
tccgcgttttctgtactatttctcccagagccgtttattttgcataggggggtgggtcataaacggagacgttttggt
ataaaatttgccccttcttccaccgttgaaaaaattggaaaatgaaccgggacaggaaaggggatttcaggataatg
gt
